# CRISPR/Cas with ribonucleoprotein complexes and transiently selected telomere vectors allows highly efficient marker-free and multiple genome editing in *Botrytis cinerea*

**DOI:** 10.1101/2020.01.20.912576

**Authors:** Thomas Leisen, Fabian Bietz, Janina Werner, Alex Wegner, Ulrich Schaffrath, David Scheuring, Felix Willmund, Andreas Mosbach, Gabriel Scalliet, Matthias Hahn

## Abstract

CRISPR/Cas has become the state-of-the-art technology for genetic manipulation in diverse organisms, enabling targeted genetic changes to be performed with unprecedented efficiency. Here we report on the first establishment of robust CRISPR/Cas editing in the important necrotrophic plant pathogen *Botrytis cinerea* based on the introduction of optimized Cas9-sgRNA ribonucleoprotein complexes (RNPs) into protoplasts. Editing yields were further improved by development of a novel strategy that combines RNP delivery with transiently stable telomeres containing vectors, which allowed temporary selection and convenient screening of marker-free editing. We demonstrate that this approach provides vastly superior editing rates compared to existing CRISPR/Cas-based methods in filamentous fungi, including the model plant pathogen *Magnaporthe oryzae*. The high performance of telomere vector-mediated coediting was demonstrated by random mutagenesis of codon 272 of the *sdhB* gene, a major determinant of resistance to succinate dehydrogenase inhibitor (SDHI) fungicides by in bulk replacement of the codon 272 with codons encoding all 20 amino acids. All exchanges were found at similar frequencies in the absence of selection but SDHI selection allowed the identification of novel amino acid substitutions which conferred differential resistance levels towards different SDHI fungicides. The increased efficiency and easy handling of RNP-based cotransformation is expected to greatly facilitate molecular research in *B. cinerea* and other fungi.

## Introduction

*Botrytis cinerea* is a plant pathogenic ascomycete which infects more than a thousand species, triggering gray mold disease which is responsible for over a billion dollars of losses in fruits, vegetables and flowers every year [1]. Due to its worldwide occurrence, great economic importance and non-specific necrotrophic lifestyle, it has been ranked as the second most important plant pathogenic fungus [2]. Control of gray mold often requires repeated treatments with fungicides, in particular under high humidity conditions, but rapid adaption and resistance development of *B. cinerea* has dramatically reduced their efficiency worldwide in many cultures, for example in strawberry fields [3]. After germination of a conidium on the plant surface, the fungus penetrates and invades the host, rapidly killing plant cells by releasing a complex mixture of cell wall degrading enzymes, phytotoxic metabolites and proteins, and by tissue acidification [4, 5]. How host cell death is induced is not fully understood, but the invading hyphae seem to trigger the hypersensitive response, a plant-specific type of apoptosis linked to strong defence reactions [6, 7]. Furthermore, *B. cinerea* releases small RNAs (sRNAs) that can suppress the expression of defence-related genes in its host plants [8]. As a countermeasure, plants also release sRNAs aimed to suppress fungal virulence [9]. To facilitate access to genes or non-coding RNA loci that are important for pathogenesis, a gapless genome sequence of *B. cinerea* has been published recently [10]. Considerable efforts have been made to generate tools for the genetic manipulation of *B. cinerea*. *Agrobacterium*-mediated and protoplast-based transformation have been developed [11–13], and several vectors are available which facilitate the generation of mutants and strains expressing fluorescently tagged proteins for cytological studies[14]. Nevertheless, the generation of mutants remains time-consuming, partly because of the multinuclear nature of *B. cinerea*, which requires several rounds of sub-cultivation to achieve homokaryosis. Furthermore, the generation of multiple knock-out mutants is hampered by the lack of marker recycling systems for serial gene replacements, as described in some filamentous fungi [15].

The application of the clustered regularly interspaced short palindromic repeats (CRISPR)-associated RNA-guided Cas9 endonuclease activity has revolutionized genome editing and greatly facilitated the genetic manipulation in a wide range of species [16]. CRISPR/Cas is based on the introduction of double stranded breaks by the Cas9 endonuclease in the genome of an organism. Cas9 targeting occurs by complementary sequences of a single guide RNA (sgRNA), which directs the endonuclease to a genomic target sequence via a 20 bp homology region [17–19]. The sequence requirement, for Cas9 from *Streptococcus pyogenes*, is the presence of the so-called protospacer adjacent motif (PAM), a triplet NGG located immediately 3′ of the target[20]. The breaks are then repaired by non-homologous DNA end joining (NHEJ) or, if a repair template (RT) DNA homologous to sequences flanking the break is provided, by homologous recombination (HR), which allows the generation of specific edits in the genome.

CRISPR/Cas has been successfully applied in various fungal species using different strategies to deliver Cas9 and the sgRNA [21]. In most cases, codon optimized versions of Cas9 encoding genes were introduced by stable chromosomal integration or transiently via plasmids. To achieve robust expression and efficient nuclear targeting of Cas9, strong fungal promoters, codon-optimized genes and suitable nuclear localization signals fused to the protein are beneficial. Delivery of sgRNA can be achieved either via plasmids or by *in vitro* synthesized sgRNA. More recently, transformation with Cas9-sgRNA ribonucleoprotein (RNP) complexes has been successfully applied in selected fungi [22–24].

In this study, we show that CRISPR/Cas-based genome editing is highly efficient in *B. cinerea* when Cas9-sgRNA RNPs are introduced into protoplasts. By using *Bos1* as a selectable marker for gene knockouts, high frequencies of edits via NHEJ and HR were achieved. With RT containing only 60 bp homology flanks, >90% targeting efficiency was observed. Taking advantage of a transiently selectable telomere vector and high cotransformation rates of CRISPR/Cas constructs, a highly efficient marker-free editing strategy was developed, yielding up to thousands of edited transformants per transformation. The power of this approach, which was verified also for *Magnaporthe oryzae*, was demonstrated by random *in vivo* mutagenesis of a resistance-associated codon in a fungicide target gene, and application which was not possible before in filamentous fungi.

## Results

### Establishment and characterization of CRISPR/Cas editing in *B. cinerea*

To achieve strong expression and robust nuclear localization of Cas9, we tested Cas9 constructs with different nuclear localization signals (NLS) using *B. cinerea* transformants expressing a GFP-tagged synthetic Cas9 gene adapted to the low GC content of *B. cinerea* [25]. *B. cinerea* transformants expressing Cas9-GFP with a single C-terminal SV40 T antigen NLS, or with two N- and C-terminal SV40 NLS, both resulted in fluorescence distributed between cytoplasm and nuclei (Fig. 1A-B). In contrast, four tandem copies of SV40 NLS (SV40^x4^) and a duplicated NLS of the nuclear StuA protein (Stu^x2^) effectively directed Cas9 into nuclei (Fig. 1C-D).

**Fig. 1.**
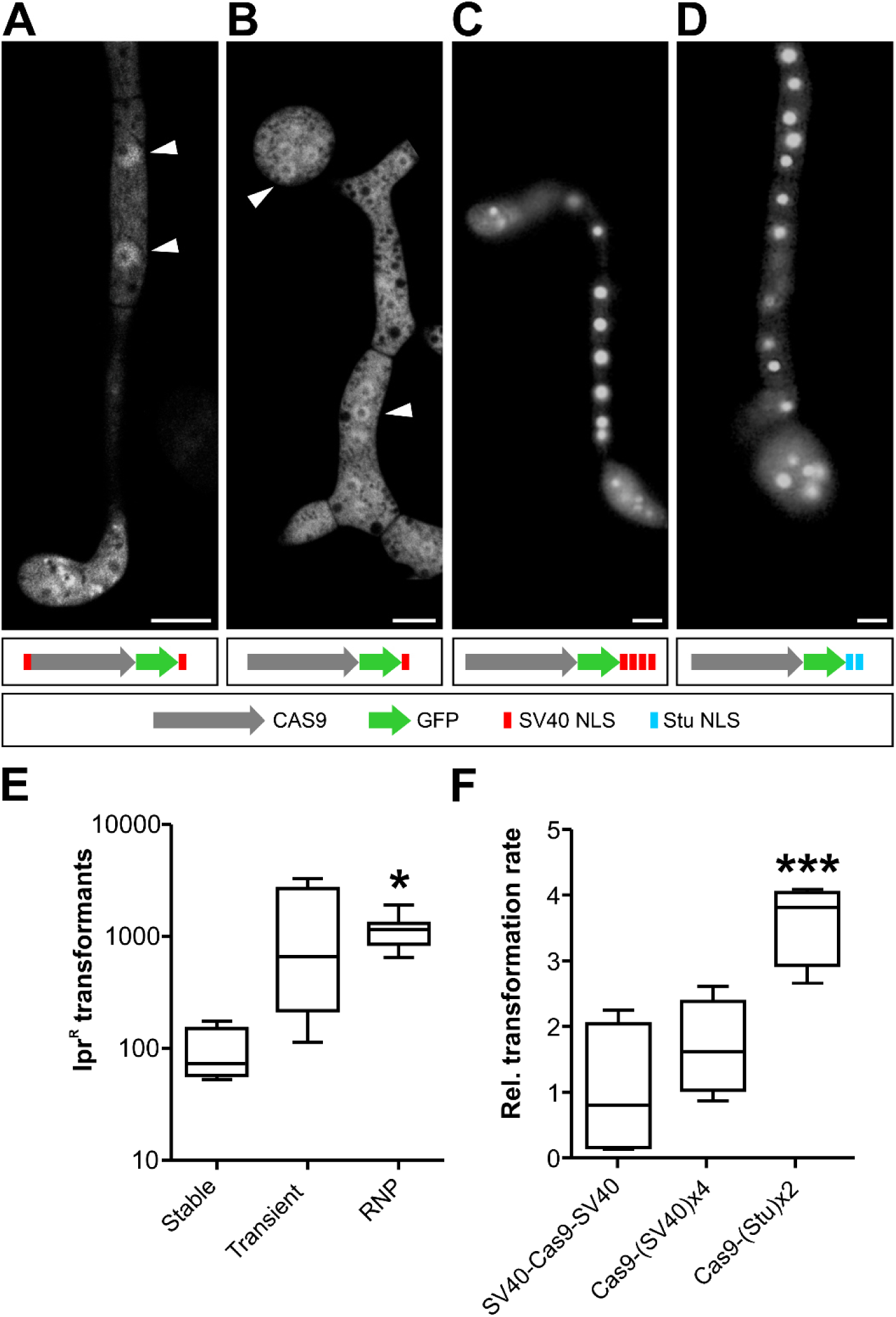
Optimization of Cas9 nuclear targeting and delivery into *B. cinerea* protoplasts. (A-D) Subcellular localization of genetically delivered Cas9-GFP constructs fused to different NLS. Fluorescence microscopy images of 18 h old germlings on glass slides. Arrowheads depict nuclei. Only in c and d, fluorescence is concentrated in the nuclei. Scale bars: 5 µm. (E) Transformation rates (NHEJ-mediated, Ipr^R^ *Bos1* k.o. transformants) obtained in *B. cinerea* with different Cas9 delivery strategies. Cas9 was expressed from a chromosomally integrated gene (Cas9-SV40^x4^-NLS; stable), transiently from a gene on a telomere vector (Cas9-GFP-SV40^x4^-NLS; transient) or added as a protein (Cas9-Stu^x2^-NLS; RNP) together with Bos1-T2 sgRNA to *B. cinerea* protoplasts. The *p* values by one-way ANOVA followed by Tuckey’s multiple comparisons post hoc test are indicated. \**p* ≤ 0.05; stable (n=4), transient (n=4), RNP (n=11). (F) Comparison of different NLS arrangements on genome editing efficiency of Cas9-sgRNA RNPs targeting *Bos1*. Values are relative to transformation rate with SV40-Cas9-SV40. In (E) and (F), no Ipr^R^ colonies were obtained without Cas9. The *p* values by one-way ANOVA followed by Dunnett’s multiple comparisons post hoc test are indicated. \*\*\**p* value ≤ 0.001; n=4.

We next tested which strategy was best suited for Cas9 delivery into *B. cinerea* protoplasts. For stable expression, a construct constitutively expressing Cas9-SV40^x4^ was first integrated into the *niaD* region of the genome. For transient expression, Cas9-GFP-Stu^x2^ cloned into a telomere vector (see below) was transformed into wild type *B. cinereal* [26]. Expression of Cas9 was confirmed by immunoblot analysis (S1 Fig). Alternatively, purified Cas9-Stu^x2^ protein assembled with a sgRNA to a ribonucleoprotein complex (RNP) were used for transformation of wild type *B. cinerea*. CRISPR/Cas activity was evaluated by quantification of error-prone repair via NHEJ, using the *Bos1* gene as a target*. Bos1* encodes a histidine kinase that regulates high osmolarity adaptation via the mitogen activated protein kinase Sak1 [27], which allows for robust positive selection of *Bos1* null mutants which have been shown to be resistant against the fungicides iprodione (Ipr) and fludioxonil (Fld) [28, 29]. With transiently expressed Cas9-GFP-Stu^x2^ and with Cas9-Stu^x2^ RNPs, high numbers of transformants were obtained, whereas stably expressed Cas9-SV40^x4^ yielded significantly less colonies (Fig. 1E). All transformants tested were both Ipr^R^ and Fld^R^, and failed to produce sporulating aerial hyphae. Compared to the wild type (WT), growth of the transformants was more strongly inhibited on media with high osmolarity, and their virulence was strongly reduced when inoculated on tomato leaves (S2 Fig). These phenotypes are consistent with those reported for *Bos1* k.o. mutants [28], and confirmed that *Bos1* was inactivated in the transformants. Due to high, reproducible transformation rates obtained, RNP-mediated transformation was used in all subsequent experiments. When recombinant Cas9 protein variants carrying different NLS were compared for their efficiency in RNP-mediated transformation, Stu^x2^ NLS was found to confer the highest *in vivo* editing activities (Fig. 1F).

To further characterize CRISPR/Cas-NHEJ editing, Cas9-Stu^x2^-NLS RNPs with different sgRNAs targeting *Bos1* between codons 344 and 372 were introduced into *B. cinerea* protoplasts. resulted in variable rates of NHEJ-induced mutations (Fig. 2 and S3 Fig). Variable editing frequencies were obtained, which correlated only weakly with *in silico* predictions and with *in vitro* cleavage assays (S3 Fig). RNP-induced *Bos1* mutations in Ipr^R^ transformants were characterized. Most of them showed insertions or deletions of only one or a few nucleotides at the cleavage sites, typical for error-prone NHEJ repair of CRISPR/Cas-induced DNA breaks. With three sgRNAs, a ‘+T’ insertion was predominant, while another sgRNA (Bos1-T7) yielded mostly three types of 9 bp deletions (Fig. 2). Insertions >1bp where found in only 19% of the transformants analyzed. Among 14 insertions of 15-164 bp, three contained *Bos1* DNA derived from sequences close to the sgRNA target sites, and one contained mitochondrial DNA (S4 Fig). The majority of insertions were derived from the scaffold used for sgRNA synthesis, which had apparently resisted the DNase treatment. Several more complex insertions were observed that involved amplification of neighboring *Bos1* sequences (S4 Fig). Taken together, our results show that with selected sgRNAs, error-prone NHEJ-mediated repair in *B. cinerea* results in remarkably uniform mutation patterns.

**Fig 2.**
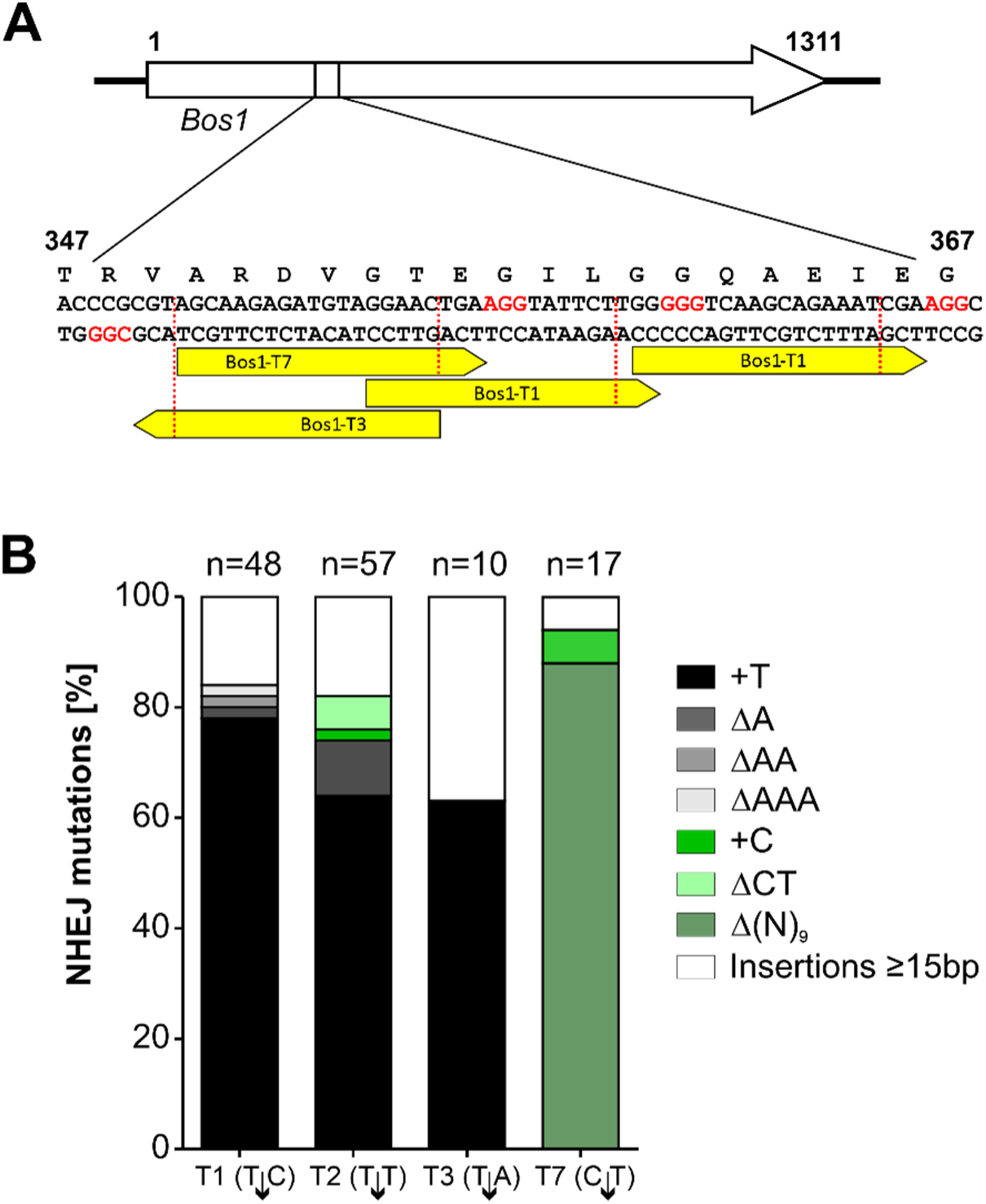
NHEJ-mediated mutations induced in *Bos1* by RNP mediated genome editing. (A) Positions of the sgRNAs targeting *Bos1*. (B) Distribution of mutations detected in iprodione resistant transformants. Note that sgRNAs introducing T↓N cleavage sites resulted in a majority of ‘+T’ insertions.

### Targeted CRISPR/Cas-mediated editing

To generate targeted *B. cinerea* insertion mutants, and to compare NHEJ- and HR-editing frequencies, a fenhexamid resistance cassette (Fen^R^) [30] flanked by 1 kb *Bos1* sequences was delivered as a repair template (RT) in addition to the RNP targeting *Bos1* into protoplasts. Transformants were selected for Fen^R^ or Ipr^R^. Regardless whether the RT was provided as circular plasmid or as PCR product, transformation rates were lower with selection for Fen^R^ than for Ipr^R^. Almost all Fen^R^ transformants were also Ipr^R^, indicating highly efficient HR-mediated integration of the Fen^R^ cassette into *Bos1*. In contrast, only 22-39% of Ipr^R^ transformants were Fen^R^, indicating a 2.5 to 5-fold higher frequency of NHEJ compared to HR in this experiment (S5 Fig).

Conventional gene targeting in filamentous fungi requires resistance cassettes with ≥0.5-1 kb flanking homology regions. A major advantage of HR using CRISPR/Cas is that dsDNA repair can also be achieved using short RT homology flanks [23, 31–33]. To test this for *B. cinerea*, Fen^R^ cassettes with *Bos1* homology flanks adjacent to the PAM sequence, ranging from 0 to 60 bp, were generated as RT. When delivered as Cas9-RNPs, the numbers of Fen^R^ transformants increased with increasing flank sizes, reaching highest values with 60 bp flanks (Fig. 3A and 3B). All Fen^R^ transformants tested were Ipr^R^, indicating correct targeting of *Bos1*. Remarkably, even 66% of the transformants obtained with a Fen^R^ cassette lacking homology flanks were also Ipr^R^. Sequencing confirmed that the cassette had integrated via NHEJ into the cleavage site in *Bos1*. When the Fen^R^ cassettes were delivered without RNP, only few Fen^R^ transformants were obtained, and none of them were Ipr^R^, indicating integrations outside *Bos1* (Fig. 3B). When the 60 bp flanks of the RT were separated by 1 kb each from the PAM site to generate a *Bos1* deletion instead of an insertion, similar transformation efficiencies were obtained (Fig. 3C and 3D). Thus, CRISPR/Cas allows the use of short homology flanks in a flexible way for highly efficient gene targeting.

**Fig. 3.**
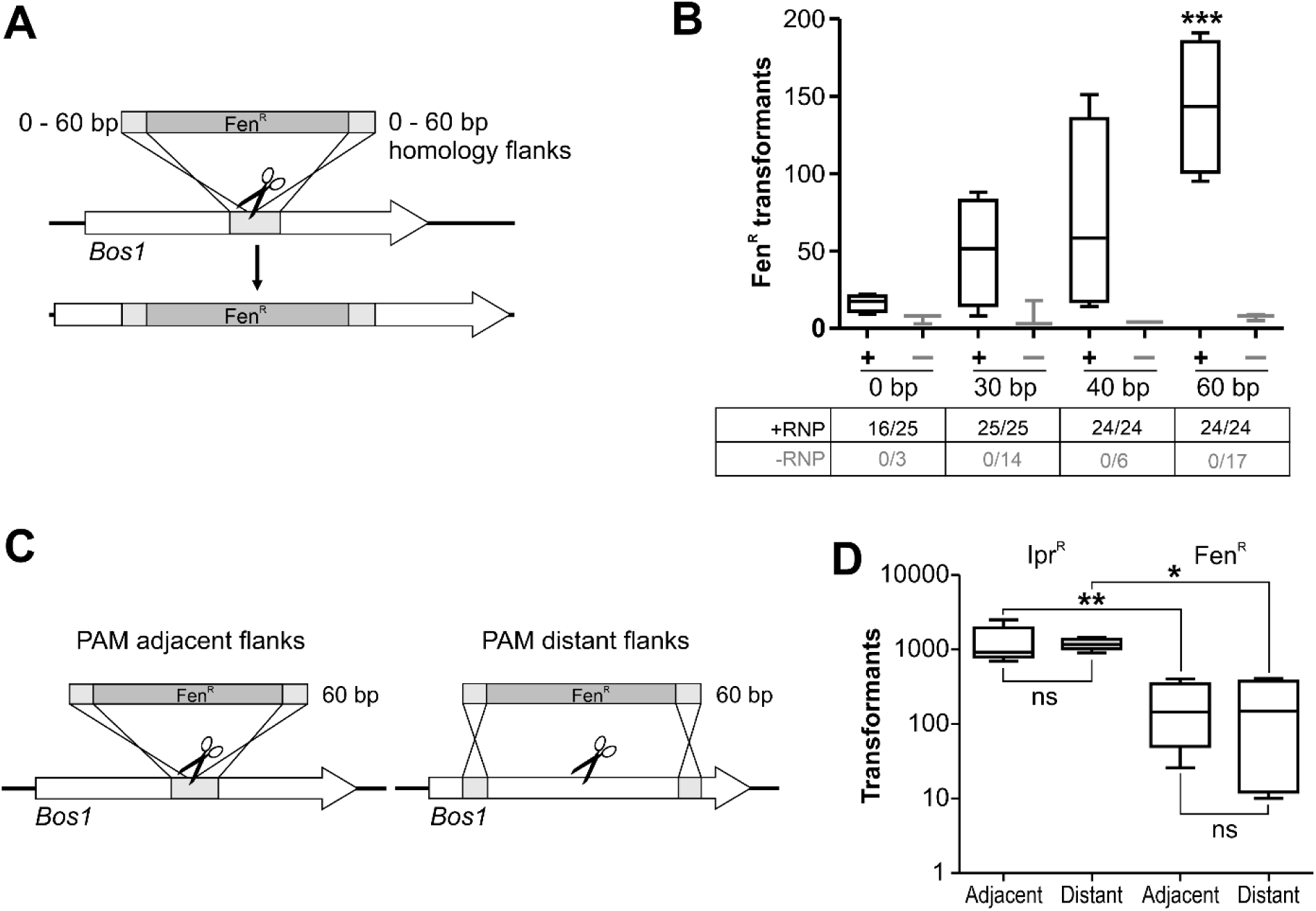
Efficiency of CRISPR/Cas editing of *Bos1* using repair templates with short homology flanks. (A) Experimental scheme. (B) Results of transformations with RNP (black lines) and without RNP (gray lines) for RT with flank sizes of 0 to 60 bp. (+RNP: n=4; RNP: n=3). The numbers below show fractions of Fen^R^ transformants being Ipr^R^, indicating targeting efficiencies. The p values by one-way ANOVA followed by Dunnetts’s multiple comparisons (control: 0 bp) post hoc test are indicated. ***p ≤ 0.001. (C) Scheme of *Bos1* targeting with different placement of 60 bp homology flanks of RT, resulting in insertion (left) or 2 kb deletion (right). (E) Results of transformations with RNP and two types of RT as shown in (C). n=5 (Ipr^R^); n=4 (Fen^R^). The *p* values by one-way ANOVA followed by Tukey’s multiple comparisons post hoc test are indicated. *p ≤ 0.05; ** p ≤ 0,01.

To exploit the efficiency of CRISPR/Cas, co-targeting of two genes encoding key enzymes for biosynthesis of the phytotoxins botrydial (*bot2*) and botcinic acid (*boa6*) was tested. The role of these toxins for *B. cinerea* is not yet completely clear. Whereas single *bot2* and *boa6* knockout mutants did not reveal a decreased pathogenicity, double mutants were found to be impaired in growth and virulence [34]. Cas9-RNPs and RTs with 60 bp flanks targeting *bot2* (using a Fen^R^ cassette) and *boa6* (using a cyprodinil (Cyp^R^) cassette) were generated. In two transformations, 39 and 47 Fen^R^ colonies, and 16 and 14 Cyp^R^ colonies, respectively, were obtained (S6A Fig). Of 70 Fen^R^ transformants tested, 49 were Cyp^R^, indicating successful coediting. PCR-based DNA analysis of 20 Fen^R^ Cyp^R^ transformants revealed 15 transformants as *boa6* k.o., four as *bot2* k.o., and three as *boa6bot2* double k.o., two of which could be purified to homokaryosis (S6B Fig). Thus, double knock-outs can be obtained with Cas-RNPs with high frequency. Phenotypical characterization of the double mutants revealed no significant differences to the WT in their vegetative growth and infection (S6C and S6D Fig). This indicated that the phytotoxins botrydial and botcinic acid are not important for *B. cinerea* to infect tomato leaves.

### Resistance marker shuttling, a simple strategy for marker-free editing

To generate precise and multiple changes in the genome, marker-free editing is required. Two marker-free mutagenesis strategies were developed, both exploiting the high efficiency of cotransformation, namely that two or more DNA constructs are taken up by fungal cells with much higher frequencies than expected from single transformation rates. The first strategy, called resistance marker shuttling, is based on the integration of an RT into a non-essential genomic locus in exchange for an existing resistance cassette with identical promoter and terminator sequences which serve as homology flanks. To test for marker exchange, a *B. cinerea* strain carrying a nourseothricin (Nat^R^) cassette in the *xyn11A* locus [35] was transformed with Cas9-RNP and a Fen^R^ RT which shared the promoter (*PtrpC*) and the terminator (*TniaD*) sequences with the targeted *nat1* gene as homology flanks (Fig. 4A). Transformations resulted in several hundred Fen^R^ colonies, and the majority of them had lost Nat^R^ as expected for a marker exchange. When *Bos1*-RNP was cotransformed, similar numbers of Fen^R^ transformants were obtained, and 56-74% of them were also Ipr^R^, demonstrating a high rate of NHEJ coediting. No marker exchange was observed when the Fen^R^ RT was transformed without Cas9-RNP as negative control (Fig. 4B). To test the stability of both resistance markers in the Fen^R^ Ipr^R^ double transformants, each ten of them were transferred three times to ME agar plates containing only Fen or Ipr. All transformants treated this way retained the non-selected resistance, indicating that coediting had occurred in the same nuclei of the transformed protoplasts. The resulting transformants could be used for another round of marker shuttling, now targeting the Fen^R^ resistance cassette.

**Fig. 4.**
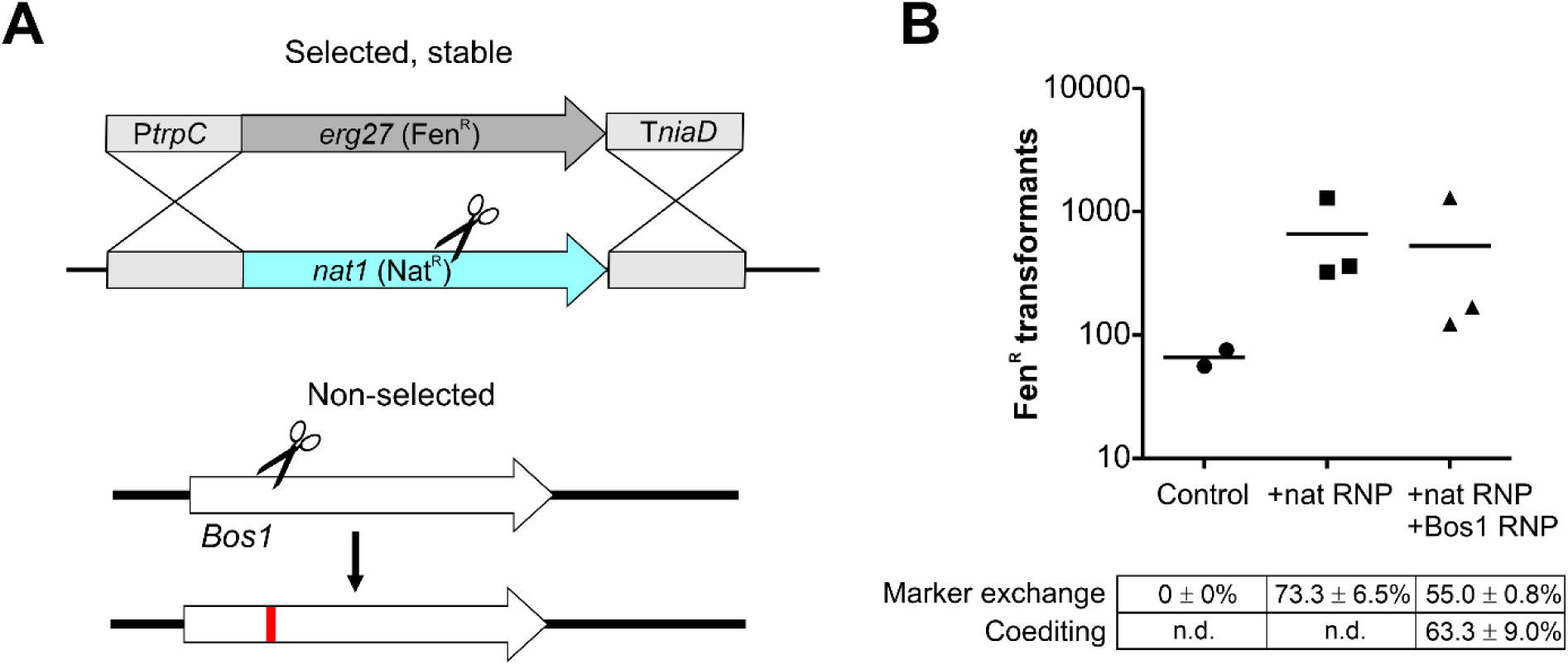
Application of marker exchange for non-selected CRISPR/Cas k.o. mutagenesis of *Bos1* via NHEJ. (A) Experimental scheme. (B) Results of transformation of *B. cinerea xyn11A*-Nat^R^ with Cas9-RNP targeting *nat* using Fen^R^ RT, with or without Cas9-RNP targeting *Bos1*, as shown in (B). Control: Fen^R^ RT transformed without Cas9-RNP.

### Use of transiently selected telomere vectors for completely marker-free coediting

Previous studies have shown that plasmids containing a pair of telomeres (pTEL) can be efficiently transformed into filamentous fungi and replicate there autonomously as centromere-free minichromosomes, but are rapidly lost in the absence of selective pressure [26]. Based on these properties, a pTEL-mediated strategy for marker-free CRISPR/Cas coediting was developed, involving the following steps (Fig. 5A and 5B): i) cotransformation of pTEL and Cas9-RNP (with or without RT) into *B. cinerea*; ii) selection for pTEL-encoded resistance; iii) identification of transformants with desired coediting events; iv) purification of the transformants by transfers on selective media until homokaryosis is confirmed; v) elimination of pTEL by transfers on nonselective media. This strategy was tested first with pTEL-Fen and Cas9-RNP targeting *Bos1* to generate k.o. mutants via NHEJ. Compared to high transformation rates obtained with pTEL-Fen alone, only few Fen^R^ colonies were obtained with 0.5-2 µg pTEL-Fen added together with Cas9-RNP to the protoplasts. The suppression of pTEL-Fen transformation by Cas9-RNP was largely overcome by increasing the amount of pTEL-Fen in the transformation mixture up to 10 µg (Fig. 5C). When Fen^R^ transformants were transferred to Ipr containing medium, 25-53% (average 40.0 ± 11.2%) of them were Ipr^R^, which demonstrated a high rate of coediting. After two transfers on nonselective medium, 22 of 26 Ipr^R^ transformants were Fen^S^, confirming the expected loss of pTEL-Fen. Few Ipr^R^ transformants remained Fen^R^ after further passages, indicating integration of pTEL-Fen into the genome. In the next cotransformation experiment with pTEL-Fen, *sod1* encoding the major copper/zinc superoxide dismutase was targeted to generate a *sod1-GFP* knock-in fusion (Fig. 5D to 5G). Sod1 has been shown to be involved in oxidative stress tolerance and virulence of *B. cinerea* [36]. Several thousand Fen^R^ transformants were obtained in single experiments (Fig 5D). Microscopic evaluation revealed GFP fluorescence in 65.3% of the transformants resulting from coediting. Fluorescence was observed in the cytoplasm and in strongly fluorescent punctate structures tentatively identified as peroxisomes (Fig. 5E). For SOD1 of rat, an orthologue of the fungal Sod1, a localization similar to *B. cinerea* was found in the cytoplasm and in peroxisomes, due to its binding to peroxisomal protein CCS [37]. The functionality of the Sod1-GFP fusion protein was confirmed by staining for SOD activity after native gel electrophoresis of protein extracts (Fig. 5F) and by immunoblotting using GFP antibodies (Fig. 5G). Furthermore, pTEL-mediated coediting was shown to be useful for marker-free mutagenesis of *nep1* and *nep2*, two genes encoding necrosis and ethylene-inducing proteins [38]. With single targeting, >1,000 Fen^R^ transformants were obtained, and 17-23% of these contained *nep1* or *nep2* deletions, respectively, as confirmed by PCR. Co-targeting of *nep1* and *nep2* resulted in 230 transformants. Of these, 12.9% contained a *nep1* deletion and 10% a *nep2* deletion (S7 Fig), but no double transformants were detected.

**Fig. 5.**
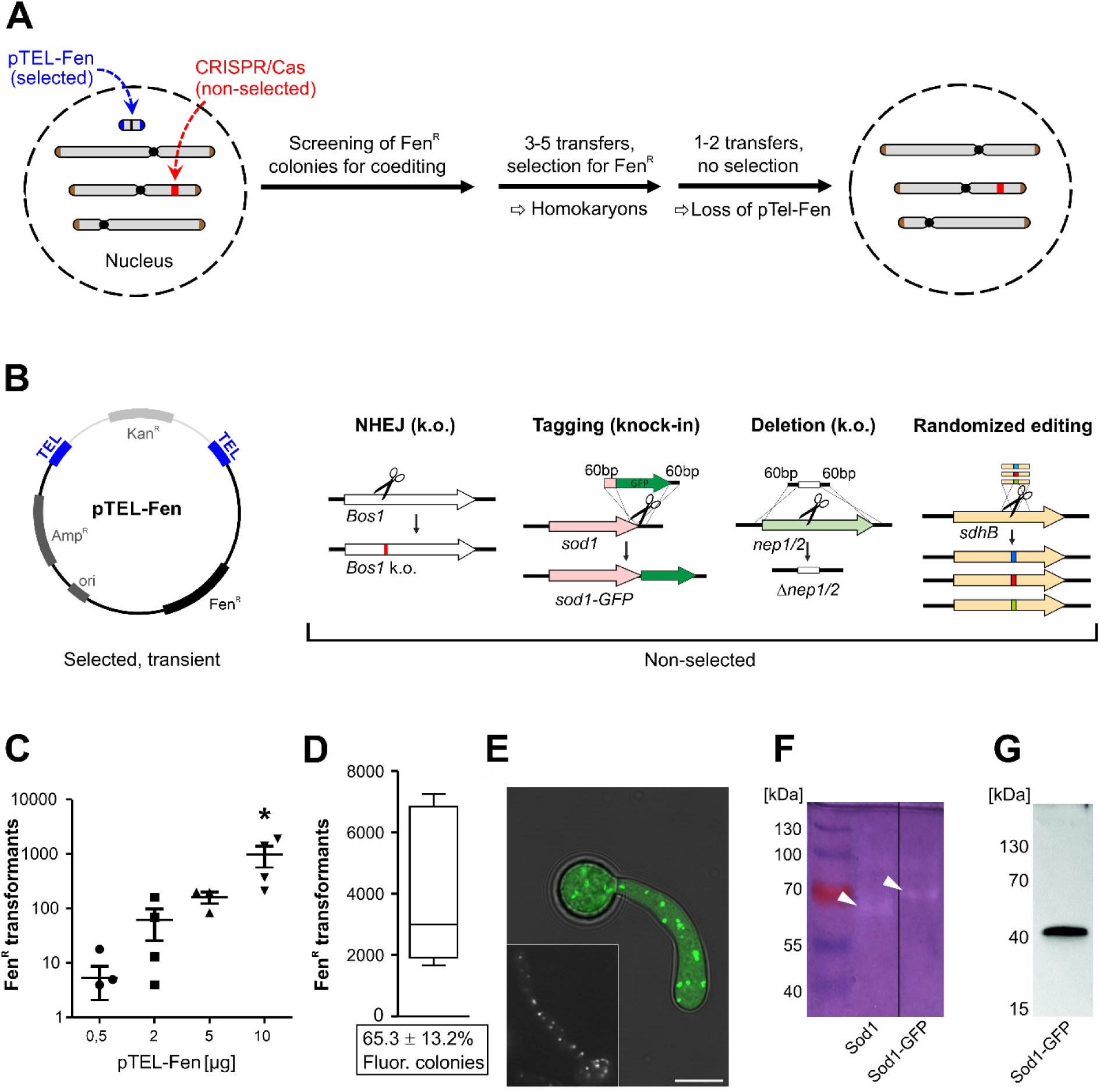
Telomere vector (pTEL)-mediated coediting for introduction of marker-free CRISPR/Cas edits into *B. cinerea*. (A) Experimental setup. (B) Applications of non-selected editing performed in this study. pTEL-Fen can be propagated in *E. coli* with selection for ampicillin (Amp^R^) and kanamycin (Kan^R^). After transformation into *B. cinerea* it is linearized to a minichromosome with telomeric ends. (C) Transformation results for pTEL-mediated *Bos1* k.o. via NHEJ, depending on the amounts of pTEL-Fen added to the protoplasts. Individual data points are shown. The *p* values by one-way ANOVA followed by Dunnetts’s multiple comparisons (control: 0.5 µg pTEL-Fen) post hoc test are indicated. *p ≤ 0.05. (D-G) Generation and characterization of a Sod1-GFP knock-in strain. (D) Transformation efficiency and frequency of fluorescent transformants (below; n=3). (E) Cytoplasmic and putative peroxisomal localization of Sod1-GFP fluorescence, as indicated by similar fluorescence pattern of a mutant expressing GFP fused to a SKL peroxisomal targeting motif [14). (F) Native gel electrophoresis of *B. cinerea* protein extracts stained for superoxide dismutase activity. Lanes showing WT (expressing Sod1) and mutant (expressing Sod1-GFP, arrowheads). (G) Immunoblot detection of Sod1-GFP with GFP antibodies.

### Telomere vector-mediated marker-free coediting also works efficiently in *Magnaporthe oryzae*

The rice blast fungus *M. oryzae* is of great economic importance and considered as the prime model pathogenic fungus [2]. It is a hemibiotroph and well-known for its ability to develop enormous turgor pressure in appressoria facilitating penetration of host cells [39]. Recently, protoplast transformation with CRISPR/Cas using RNP has been successfully applied for this fungus [23]. Marker-free editing was also demonstrated, however, rates of non-selected coediting events ranged only from 0.5 to 1.2%. Aiming to improve this rate, we first confirmed the efficacy of CRISPR/Cas with Cas9-RNP. *M*. *oryzae* strain Guy11 or Guy11ku80 (a NHEJ deficient mutant) protoplasts were transformed with Cas9-SV40^x4^ complexed with sgRNA *MoALB1* and a Hyg^R^ RT with 50 bp homology flanks. *MoALB1* encodes a polyketide synthase required for melanin biosynthesis and *alb1* mutants are easily selectable due to whitish mycelium. Depending on the amount of RT DNA and strain used, 67 to 91% of Hyg^R^ transformants had white mycelium, indicating successful inactivation of *MoALB1* (S8 Fig).

Next, we tested the suitability of the pTEL-based marker-free approach for *M. oryzae*. After establishing selection for Fen^R^, using 30 ppm fenhexamid, pTEL-Fen was transformed yielding up to 1,000 transformants per µg DNA (S1 Table). Subsequently, pTEL-Fen was cotransformed together with Cas9-sgRNA RNP targeting *MoALB1*. Among Fen^R^ transformants, 36-49% displayed white colonies in Guy11, indicating a high rate of co-editing (Fig. 6). By contrast, the rate of cotransformation in Guy11ku80 was much lower. Sequencing of *MoALB1* in three of the white Fen^R^ colonies of Guy11 revealed the presence of single base pair deletions at the cleaving site, leading to frameshifts. After two passages on non-selective medium, 12 out of 15 albino mutants were Fen^S^, as predicted from the instability of pTEL-Fen. To show that coediting with insertion of a RT into a specific locus is possible as well, pTEL-Fen, Cas9-RNP targeting the *MoPIT* gene MGG_01557 and a Hyg^R^ cassette with 50 bp flanks were cotransformed into *M. oryzae* protoplasts (S9 Fig). While 23 out of 72 Fen^R^ transformants were Hyg^R^, *MoPIT* knockouts were detected in five of these, representing a coediting rate via HR of 7%. However, no coediting was observed in *M. oryzae* Guy11ku80.

**Fig. 6.**
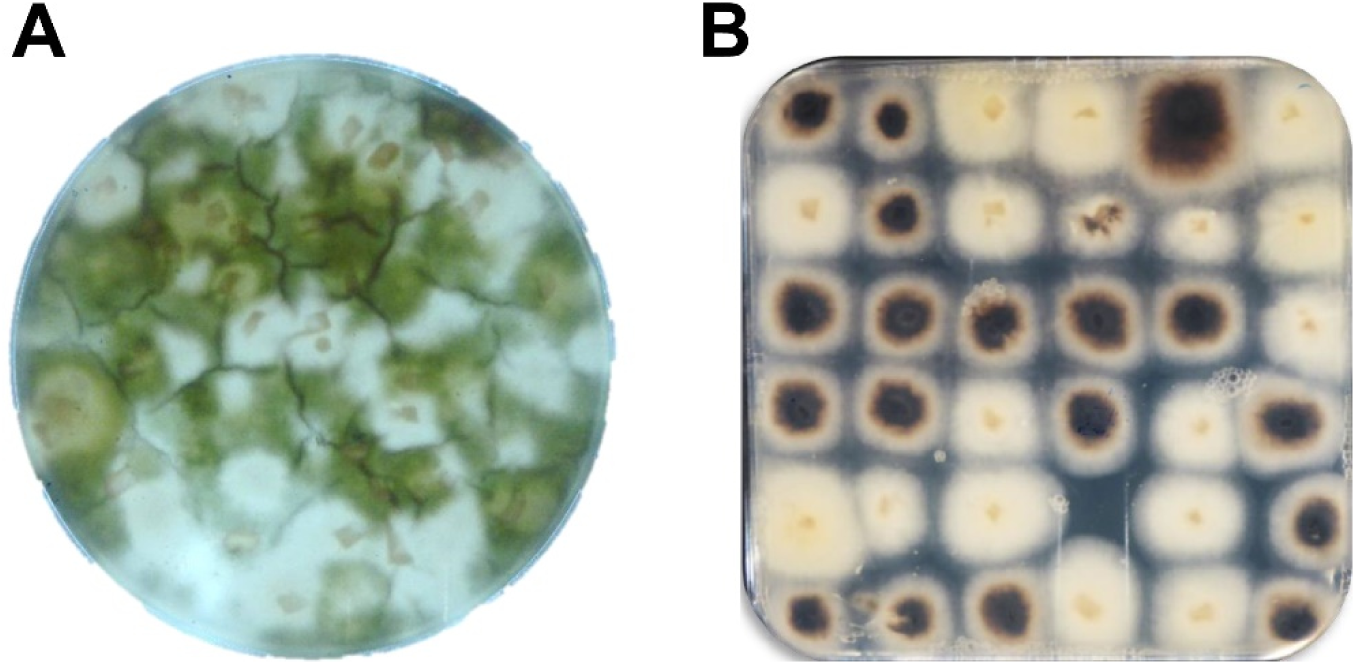
Efficient pTEL-mediated coediting via NHEJ in *Magnaporthe oryzae*. Protoplasts were cotransformed with pTEL-Fen and Cas9-ALB1-sgRNA RNP. (A) Primary selection plate containing fenhexamid. (B) Isolated transformants (transformation C, cf. S1 Table). Note white-colored mycelia of edited transformants.

### Randomized amino acid editing of a fungicide resistance codon and *in vivo* selection

Succinate dehydrogenase inhibitor (SDHI) fungicides have emerged as the fastest increasing class of fungicides for the control of plant diseases in recent years [40]. Target site mutations leading to resistance against SDHI have been described in *B. cinerea* and other fungi. Most of them are located in *sdhB* encoding the B-subunit of the succinate dehydrogenase enzyme (complex II), an essential component of mitochondrial respiration. In *B. cinerea* populations from SDHI-treated fields, *sdhB* mutations leading to H272R, H272Y, H272L, and H272V amino acid exchanges have been found [41]. While all of them confer resistance to boscalid (Bos), only H272L and H272V mutations also confer resistance to another SDHI, fluopyram (Flu) [40, 42]. To analyze the effects of all possible exchanges in codon 272 for SdhB function and resistance against SDHIs, pTEL-mediated coediting was performed to target *sdhB* with a RT mixture encoding all 20 amino acids in codon 272 (Fig. 7A). Several thousand colonies were obtained per transformation. Estimations based on PCR analysis of single transformants revealed coediting frequencies between 12.5 and 41% (S2 Table). The distribution of codons in position 272 was determined from pooled conidia of ≥6,000 Fen^R^ transformants per assay by bulked DNA isolation, followed by deep sequencing. Aliquots of pooled transformant conidia were cultivated for three days in liquid medium containing discriminatory concentrations of Bos, Flu, or the new SDHI fungicide pydiflumetofen (Pyd) [43], to select transformants with SDHI resistance. DNA of these cultures was isolated and sequenced as above. Among the edited transformants grown on SH+Fen plates, all 20 codons were represented at similar frequencies (Fig. 7B). Because in this procedure edited cells may still carry a WT copy of *sdhB* (heterokaryons), we cannot conclude yet that they all maintained full enzyme function. However, our results demonstrate that all H272 amino acids variants yield functional SdhB proteins that are not intrinsically toxic since they all similarly maintained growth and sporulation. Cultivation of the Fen^R^ transformants in SDHI-containing media followed by quantification of the alleles enabled an unbiased assessment of amino acid exchanges conferring resistance. Most conspicuous resistance mutations were H272K/V/L/R/Y for Bos, H272N/L/V/I for Flu, and H272V/L for the new SDHI Pyd (Fig. 7C-7E). Transformants with 17 different exchanges in codon 272 were isolated, purified to homokaryosis, and tested for sensitivity to the three SDHI (Fig. 7F). Overall, the EC_50_ values correlated well with their prevalence in the SDHI selected populations described above (Fig. 7c-d). Remarkably, 12 amino acids conferred high levels of resistance to boscalid (EC_50_ values >2mg l^-1^) (Fig. 7C). In contrast, only four amino acids conferred similarly high resistance levels to Flu, but five amino acids caused up to 30-fold hypersensitivity compared to WT (Fig. 7D). Pyd was about ten times more active than Bos and Flu against *B. cinerea* WT, and four amino acids conferred EC_50_ values >0.2mg l^-1^ (Fig. 7F). Only the three aliphatic amino acids leucine, valine and isoleucine provided high or intermediate resistance to all three SDHI. Remarkably, highest resistance levels were observed with isoleucine, which has never been found in resistant field isolates. Growth on selective agar media illustrated the high proportion of Bos^R^ mutants, and the lower number of mutants resistant to Flu and Pyd (Fig. 7G). Growth on rich medium and on a nutrient-limited medium with different carbon sources did not reveal significant differences between the 17 edited strains (S10 Fig), indicating no major effects of the amino acids on the fitness of the mutants during vegetative growth.

**Fig. 7.**
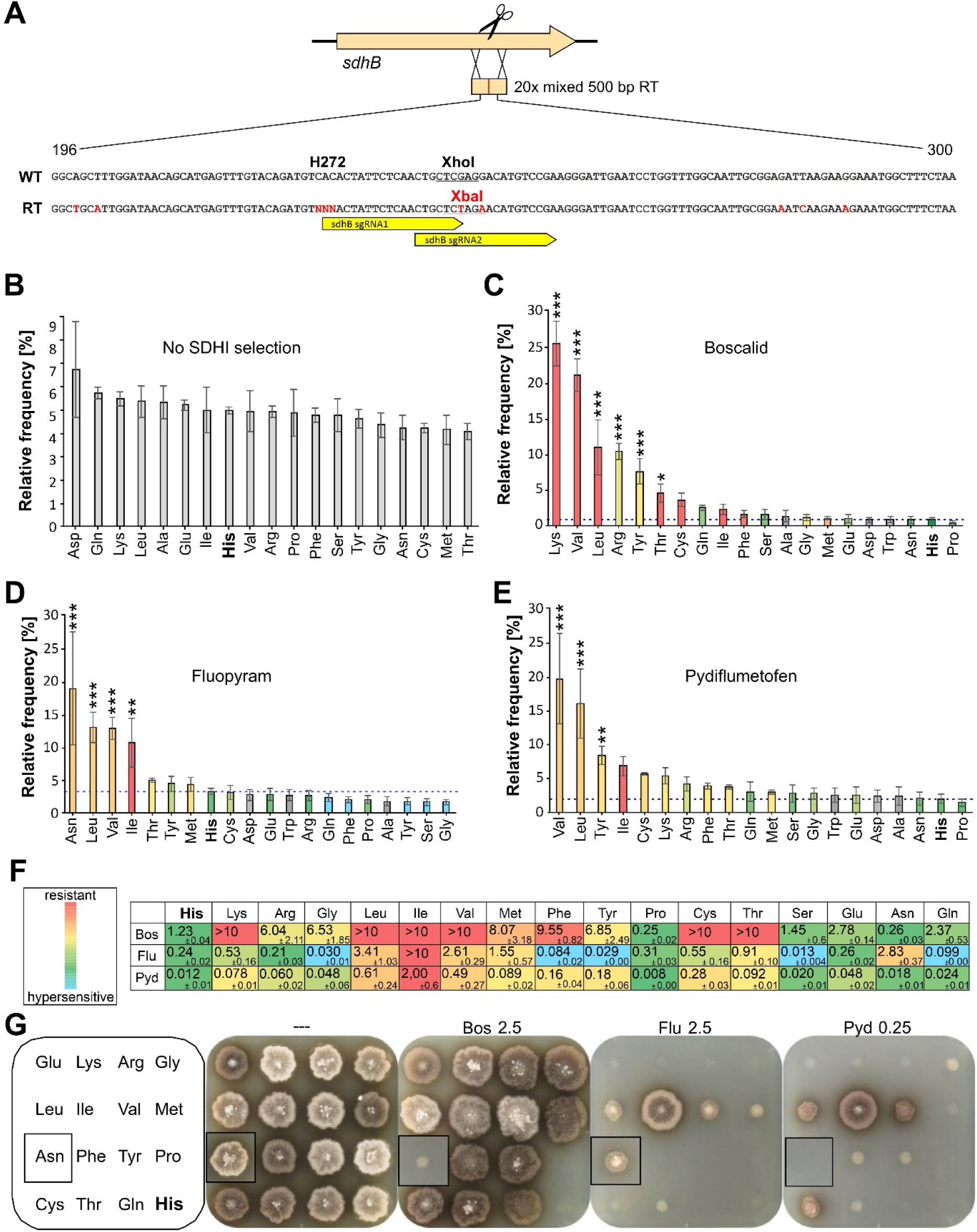
Effects of *B. cinerea sdhB* codon 272 amino acid randomization by multiple editing. (A) Schematic strategy, showing sequences of WT and repair template (RT) surrounding codon 272. Changed bases and the new restriction site in the RT are marked in red. NNN: Each of 20 codons in the RT mixture. (B-E) Frequency distribution of encoded amino acids in codon 272 of edited *B. cinerea* transformants, determined by deep sequencing of conidia obtained from primary Fen^R^ transformants without SDHI fungicide selection (B), or from cultures of transformed conidia incubated in YBA medium containing 0.25 mg l^-1^ Bos (C), 0.5 mg l^-1^ Flu (D), or 0.15 mg l^-1^ Pyd. (E) Fungicide sensitivity levels conferred by each amino acid, as determined for the corresponding mutant are indicated by colors (in (C-F)), except for the bars of amino acids for which no mutants were obtained which are shaded in gray. The *p* values by one-way ANOVA followed by Dunnetts’s multiple comparisons (control: His) post hoc test are indicated. \**p* ≤ 0.05, \*\**p* ≤ 0.01, \*\*\**p* ≤ 0.001 (n=3). (F) SDHI sensitivity (EC50 values in mg l^-1^) of individual mutants containing different amino acids in *sdhB* codon 272. (G) Growth of individual codon 272 edited mutants on YBA agar containing different SDHI as indicated, after 5 days. The inserts show colonies of an Asn mutant which were integrated into the pictures of plates with the other mutants.

## Discussion

Within a short time, CRISPR/Cas genome editing has been used for the genetic manipulation of a wide range of organisms, offering new perspectives in functional genomics. In fungi, advanced CRISPR/Cas systems have been mainly established for *Aspergillus* spp. [31, 44, 45] and *Ustilago maydis* [21, 46]. They take advantage of autonomously replicating circular plasmids, namely the *Aspergillus*-derived AMA1 plasmid and pMS7 in *U. maydis*, for the delivery of Cas9 and sgRNA. AMA1 has also been used in other fungi, including the plant pathogens *Alternaria alternata* [47] and *Fusarium fujikuroi* [48]. However, this plasmid displays only low transformation rates in *B. cinerea* (S. Fillinger, personal communication). Alternatively, a non-integrating vector with human telomeres [26, 49] has been developed in this study as a tool for coediting in *B. cinerea* and *M. oryzae*.

This is the first report of powerful use of CRISPR/Cas in *B. cinerea*. A crucial step was the generation of a fully functional, nuclear targeted Cas9. SV40 NLS has been used frequently [21], but efficient nuclear targeting of Cas9-GFP-NLS has been confirmed only in some fungi [24, 50] or optimal activity experimentally verified [48]. In *B. cinerea*, efficient Cas9 nuclear targeting was achieved with C-terminal tandem arrays of either SV40 (4x) and *stuA* (2x) NLS sequences. In most fungi, CRISPR/Cas activity was detected in pilot studies by targeting genes for the biosynthesis of melanin [23, 44]. Following the strategy reported for *Fusarium graminearum* [51], *Bos1* was established as an effective selectable marker for NHEJ- and HR-mediated mutagenesis (Fig. 1 and 3). Introduction of Cas9-sgRNA RNPs with or without a donor template yielded hundreds to thousands of edited *B. cinerea* transformants. So far, similar approaches have been rather rarely used for CRISPR/Cas genome editing in fungi [22–24]. An advantage of the use of RNP over endogenous Cas9 and sgRNA expression is the reduced probability of potential off-target mutagenic activities of Cas9, because of its limited stability in cells [52]. Furthermore, sgRNAs synthesis is performed quickly and does not require any cloning steps.

A total of 153 NHEJ repair events in the *Bos1* gene were analyzed, which is the largest number reported for filamentous fungi. Most changes were 1-2 bp indels, and for three sgRNAs inducing a T↓N cleavage by Cas9, a (+T) insertion was the dominating mutation. Although all these mutations were biased by the selection for loss of *Bos1* function (Ipr^R^), these data are in line with systematic studies of CRISPR/Cas-NHEJ mutations in human cells and yeast [53, 54] which often resulted in +1 bp insertions at the Cas9-RNP cleavage site. This rather reproducible NHEJ repair in *B. cinerea* could be exploited to introduce predictable frameshift mutations even without RT. Furthermore, we demonstrate that RTs with 60 bp homology flanks worked efficiently in *B. cinerea*, yielding >90% targeted integrations (Fig. 3). Such short flanks can be attached to a resistance cassette of choice using long PCR primers, avoiding time-consuming cloning or amplification steps which were previously required to generate the long homology flanks for conventional targeted integration.

In *B. cinerea*, cotransformation occurred with rates of up to >60%, both for different combinations of CRISPR/Cas-induced integrations (HR/HR or HR/NHEJ) and for telomere vector uptake and CRISPR/Cas events (HR or NHEJ). Cotransformation rates were found to increase with higher DNA concentrations, consistent with early reports for fungi [55]. Two novel strategies have been established for marker-free coediting. Resistance marker shuttling at a non-essential locus in combination with non-selected CRISPR/Cas events allows repeated genomic edits. High frequencies (65.3%) of marker replacement were observed in the transformants, and this approach is also applicable for other organisms. A prerequisite for successful coediting in multinuclear fungi such as *B. cinerea* is the generation of homokaryons, which requires integration of different DNA fragments into the same nuclei. In most transformants analyzed this was found to be the case, similar to previous reports for *Neurospora crassa* [56].

The most powerful approach for marker-free editing is cotransformation of pTEL vector and CRISPR constructs. Its effectiveness depends on i) high transformation efficiency of pTEL which provides the selection, ii) high rates of cotransformation/coediting of pTEL and CRISPR components, iii) highly efficient HR, and iv) elimination of pTEL after identification of the desired editing event(s), yielding edited strains without any other genomic alterations. With this approach, we reproducibly obtained hundreds to thousands of transformants, and up to >50% of them were marker-free edits. Similar results were obtained for NHEJ- and HR-induced edits, as shown for NHEJ-mediated mutagenesis of *Bos1*, RT-mediated knock-in attachment of a GFP tag to *sod1*, and deletion of *nep1* or *nep2*. Importantly, we could show that the pTEL strategy is also applicable for coediting approaches in other filamentous fungi. pTEL-Fen transformed *M. oryzae* with equal efficiency as *B. cinerea*, and coediting frequencies were 36-49% for NHEJ, and 7% for RT-mediated HR. These values clearly exceed coediting rates previously reported with integrative selected markers [23]. The lower rate of coediting in *M. oryzae* with HR is probably due to the intrinsically lower efficiency of HR compared to *B. cinerea*. This could be partially compensated by using RT with longer homology flanks. We therefore expect that cotransformation with pTEL vectors will significantly facilitate the establishment of RNP-based CRISPR/Cas coediting in many fungi, and maybe also in non-fungal microbes such as oomycetes.

The power of pTEL-mediated marker-free editing was exploited by performing an unbiased directed mutagenesis of codon 272 of *sdhB* encoding the succinate dehydrogenase B subunit, the gene in which most mutations conferring resistance against SDHI fungicides have been observed in *B. cinerea* field isolates [39]. Among sporulating transformants, edited strains with all amino acid substitutions were generated with similar frequencies, compared to only four changes detected in field isolates. Drastic differences were observed for the effects of each amino acid on the sensitivity or resistance to the three SDHI tested, which underlines the importance of the conserved histidine 272 for SDHI binding [57]. The majority of substitutions caused high levels of resistance to Bos, whereas fewer substitutions conferred similar resistance levels to Flu and Pyd. Our results are consistent with the observation that Bos^R^ resistant *B. cinerea* field isolates with H272R and H272Y substitutions were sensitive or even hypersensitive to Flu and still controllable by this SDHI [41]. Previously, phenotypic characterization of field isolates and of isogenic H272R, H272Y and H272L strains generated by conventional mutagenesis with simultaneous introduction of a resistance cassette at the *sdhB* target locus indicated that these substitutions caused fitness defects, such as aberrant growth and differentiation and reduced competitiveness [58–60]. Although our analysis of the edited strains did not include enzyme activity assays, their equal distribution upon primary selection and normal growth behavior on different media does not support this conclusion. Indeed, this might reflect the great advantage of precise marker-free genome editing in avoiding any modification of neighboring genes or their regulatory sequences by co-introduction of a nearby resistance cassette. Another benefit of our approach is that it allows the analysis of several independent mutants that have been obtained without selection of the target locus. This might obviate the need for tedious complementation experiments to verify the connection between mutations and phenotypes. We further show how selection post-mutagenesis can enable the rapid scanning of mutations conferring resistance to various SDHI fungicides. Since a vast set of target mutations and fungicides can be tested, this new capability is of major relevance for accelerated fungicide design. Interestingly, several substitutions conferring high resistance levels, such as H272I, H272C and H272T have not yet been detected in field populations and suggest that a bias prevented their appearance and propagation in nature. Fungicide resistance mutations are often caused by single nucleotide exchanges, for example the major mutations against most systemic fungicides in *B. cinerea* [3, 61], including the exchanges H272R/Y/L in SdhB [62]. An obvious explanation for their unequal occurrence is their differential effects on fungal fitness, therefore mutations resulting in minimal fitness costs are most likely to occur [63]. Our data, showing similar SDHI resistance and fitness levels caused by hitherto unknown substitutions seem to indicate that fungicide resistance development in field populations is also limited by the number and probability of mutations required to change one codon to another [64].

The high yield of telomere vector-mediated coediting in combination with RNP-CRISPR/Cas opens the door to advanced genome editing applications with *B. cinerea* and other fungi, such as large-scale mutagenesis and gene tagging projects. Approaches similar to mutagenesis of *sdhB* codon 272 are now possible for *in vivo* selection and structure-function analysis of proteins, such as those involved in fungicide resistance, host invasion or any other functions of interest.

## Materials and Methods

### Fungi

*Botrytis cinerea* B05.10 was used as WT strain in this study. For demonstration of CRISPR/Cas-assisted marker replacement, a *B. cinerea* B05.10 derivative, containing a Nat^R^ cassette (*PtrpC-nat-TniaD*) integrated in *xyn11A* [35] was used. Cultivation of *B. cinerea* and infection tests were performed as described [5]. Guy11 was used as *Magnaporthe oryzae* WT strain. A NHEJ-deficient *M. oryzae* mutant, Guy11ku80, was kindly provided by A. Foster.

### DNA constructs for transformations

All oligonucleotides used are listed in Supplementary Table 3. Sequences of plasmids marked with * are provided in Supplementary file 1. Derivates of the telomere vector pFAC1 [26] were constructed as following: pFAC1 was digested with BglII/NheI, and the vector fragment ligated with a synthetic linker made by annealing of oligonucleotides pFAC1-del1/pFAC1-del2, resulting in pFB2N*, carrying a hygromycin resistance cassette. For telomere vector-mediated coediting, a truncated version of pFB2N carrying a fenhexamid resistance marker was generated, called pTEL-Fen*. A codon optimized version of the *Streptococcus pyogenes cas9* gene for expression in *B. cinerea*, under the control of *oliC* promoter from *A. nidulans*, was synthesized by Genewiz (South Plainfield, NJ, USA). To generate a stable Cas9 expressing *B. cinerea* strain, a nourseothricin resistance cassette consisting of *A. nidulans trpC* promoter (Ptrpc), *nat* gene and *B. cinerea gluc* terminator (Tgluc) [14] was integrated next to the *cas9* gene, and homology flanks for targeted integration of the construct into *niaD* encoding nitrate reductase were added. For efficient nuclear localization of Cas9, a synthetic sequence encoding four copies of the SV40 T antigen NLS (SV40^x4^) was C-terminally attached to the *cas9* coding sequence, resulting in pUC-BcCas-SV40x4_nat_niaD*. To test different NLS arrangements for their efficiency to target Cas9 into nuclei of *B. cinerea*, Cas9 was fused to GFP codon-optimized for *B. cinerea* (from pNAH-OGG [14]) and the following NLS sequences C-terminally attached: Single copy SV40, SV40^x4^, Stu^x2^ (a tandem duplicated NLS of Bcin04g00280 encoding a homologue of the *A. nidulans* nuclear protein StuA [47]), and SV40^x2^ (each one N- and C-terminal SV40). For transient expression of Cas9-GFP, pFB2N was first truncated by digestion with BlpI/SphI, followed by ligation with the annealed oligonucleotides FB108/ FB109, resulting in pFB2N_BlpI_MreI. This plasmid was digested with BlpI/MreI and ligated with fragments containing Cas9-GFP-NLS, resulting in pTEL-BcCas9GFP-NLS-SV40x4* and pTEL-BcCas9GFP-NLS-Stux2*.

To generate a RT with 1 kb *Bos1* homology flanks and a fenhexamid resistance cassette, pBS-KS(-) was digested with EcoRV and combined by Gibson assembly with two adjacent 1 kb *Bos1* homology flanks, using primers TL29 pBS_ol_bos 3.REV/ TL30 pBS_ol_bos 3.FOR and TL31 pBS_ol_bos 1.FOR/ TL32 pBS_ol_bos 1.REV, and a fenhexamid resistance cassette amplified from pNDF-OCT [30] with primers TL33 Fen_ol_bos 2.FOR/TL34 Fen_ol_bos 2.REV. From the resulting plasmid (pBS_Bos1_KO_Fen), *Bos1* RT with short homology flanks were amplified with the following primers: TL37_Fen_fw/ TL38_Fen_rev (0 bp); TL65_Bos1_Fen30_fw/ TL66_Bos1_Fen30_rev (30 bp), TL67_Bos1_Fen40_fw/ TL68_Bos1_Fen40_rev (40 bp); TL69_Bos1_Fen60_fw/ TL70_Bos1_Fen60_rev (60 bp). A RT with 60 bp *Bos1* homology flanks at 1 kb distance from the cleavage site was amplified from pTel-Fen using primers TL113 60bp Bos1 PD FW/ TL114 60bp Bos1 PD RV. For generation of *boa6* k.o. mutants, a newly designed cyprodinil resistance cassette was used (Leisen et al., unpublished).

### Expression of Cas9 protein with *B. cinerea* optimized NLS

SV40^x4^ and Stu^x2^ NLS were fused to the 3’-terminus of *Streptococcus pyogenes* Cas9 (*E. coli* codon optimized) and cloned into pET24a. The resulting plasmids, pET24a_Cas9-SV40x4-NLS-His* and pET24a_Cas9-Stux2-NLS-His* were used to express these Cas9 derivatives in *E. coli* BL21(DE3) at 20°C in autoinduction medium. Cells were harvested and ca. 10 g of cell paste resuspended in 50 ml extraction buffer (20 mM HEPES, 25 mM imidazole, 500 mM NaCl, 0.5 mM TCEP, pH 8) by stirring for 40 min. Cells were lysed using a Cell Disruptor (Constant Systems Limited, Daventry, UK) at 20,000 psi, and the lysate clarified by centrifugation at 20,000 rpm in a fixed-angle rotor for 30 min, 4°C. The lysate was applied to a 5 ml HisTrap FF column equilibrated in extraction buffer. Bound protein was eluted with 3.5 column volumes of elution buffer (20 mM HEPES, 500 mM imidazole, 500 mM NaCl, pH8, 0.5 mM TCEP). The eluate was loaded onto a GE 26/60 S200 SEC column equilibrated in 20 mM HEPES, pH8, 0.5 mM TCEP. Fractions containing the target protein were pooled and 20 % (v/v) glycerol was added. The solution was concentrated using a 10 kDa Vivaspin column. Aliquots were frozen in liquid nitrogen and stored at -80°C until use. Functionality of *in vitro* assembled Cas9-sgRNA complexes was tested by *in vitro* cleavage of target DNA as described [65].

### Synthesis of sgRNA and RNP formation

Selection of appropriate sgRNAs was carried out with the help of the sgRNA design tool of the Broad Institute (https://portals.broadinstitute.org/gpp/public/analysis-tools/sgrna-design). Oligonucleotides for synthesis of sgRNAs are listed in Supplementary Table 3. DNA template preparation was performed by annealing 10 µmol each of constant sgRNA oligonucleotide (TL147_gRNA rev) and protospacer specific oligonucleotide in 10 µl in a thermocycler (95°C for 5 min, from 95°C to 85°C at 2°C sec^-1^, from 85°C to 25°C at 0.1°C^-1^), followed by fill-in with T4 DNA polymerase (New England Biolabs, Beverly, MA, USA), by adding to the annealing mix 2.5 µl 10 mM dNTPs, 2µl 10x NEB buffer 2.1 (50 mM NaCl, 10 mM Tris-HCl, 10 mM MgCl_2_, 100 µg ml^-1^ BSA, pH 7.9), 5 µl water and 0.5 µl enzyme, and incubation for 20 min at 12°C and column purification. Subsequently, sgRNA synthesis was performed using the HiScribe™ T7 High Yield RNA Synthesis Kit (NEB), and purified using the RNA Clean & Concentrator-25 kit (Zymo Research, Orange, CA, USA). Cas9-NLS, containing N- and C-terminal SV40 NLS, was purchased from NEB. For RNP formation, 6 µg Cas9 was incubated in cleavage buffer (20mM HEPES, pH 7.5, 100 mM KCl, 5% glycerol, 1 mM dithiothreitol, 0.5 mM EDTA, pH 8.0, 2 mM MgCl_2_) with 2 µg sgRNA for 30 min at 37°C.

### Transformation of *B. cinerea*

Transformation was performed based on a published protocol[5] as following: 10^8^ conidia harvested from sporulating malt extract (ME: 10 g/l malt extract, 4 g/l glucose, 4 g/l yeast extract, pH 5.5) agar plates were added to 100 ml ME medium and shaken at 180 rpm for ca. 18 h (20-22°C) in a 250 ml flask. The germlings was transferred into 50 ml conical tubes and centrifuged (8 min, 1,000 g) in a swing-out rotor. The combined pellets (fresh weight should be >3 g) were resuspended and washed two times with 40 ml KCl buffer (0.6 M KCl, 100 mM sodium phosphate pH 5.8; centrifugation for 5 min, 1,000 g), and the germlings resuspended in 20 ml KCl buffer containing 1% Glucanex (Sigma Aldrich, St Louis, MO, USA; L1412) and 0.1 % Yatalase (Takara, T017), and incubated on a 3D rotary shaker at 60 rpm for 60-90 min at 28°C until ca. 10^8^ protoplasts had been formed. Protoplasts were filtered through a sterile nylon mesh (30 µm pore size) into a 50 ml conical tube containing 10 ml ice-cold TMS buffer (1 M sorbitol, 10 mM MOPS, pH 6.3). After addition of another 30-40 ml ice-cold TMS buffer, the suspension was centrifuged (5 min, 1500 g, 4°C), and the protoplast pellet resuspended in 1-2 ml TMSC buffer (TMS + 50 mM CaCl2, 0°C, dependent on the desired protoplast concentration. To 0.5 x10^6^ to 2x10^7^ protoplasts in 100 µl TMSC, the Cas9/sgRNA ribonucleoprotein (RNP) complex (6 µg Cas9, 2 µg sgRNA; pre-complexed for 30 min at 37°C) and up to 10 µg donor template DNA were added in 60 µl Tris-CaCl_2_ buffer (10 mM Tris-HCl, 1 mM EDTA, 40 mM CaCl_2_, pH 6.3). After 5 min incubation on ice, 160 µl of PEG solution (0.6 g ml^-1^ PEG 3350, 1 M sorbitol, 10 mM MOPS, pH 6.3; pre-heated to 60°C, mixed, and allowed to cool down to 30-40°C) was added, mixed gently, and incubated for 20 min at room temperature. 680 µl of TMSC buffer was added, the sample was centrifuged (5 min, 1,500 g in a swing-out rotor), the supernatant removed, and protoplasts suspended in 200 µl TMSC. Protoplasts were transferred into 50 ml liquid (42°C) SH agar (0.6 M sucrose, 5 mM Tris-HCl pH 6.5, 1 mM (NH_4_)H_2_PO_4_, 9 g l^-1^ bacto agar, Difco) and poured into two Petri dishes. For transformation with pTEL-Fen, up to 10 µg plasmid DNA was used. For selection of transformants, 30 mg l^-1^ nourseothricin (Nat), 1 mg l^-1^ fenhexamid (Fen), 4 mg l^-1^ iprodione (Ipr), or mg l^-1^ fludioxonil (Fld) were added. Positive colonies were transferred onto ME agar plates or onto plates containing the same concentrations of selective agents. Transformants were subcultured on selective media and purified by three to five rounds of single spore isolation. Genomic DNA was isolated as described previously [66].

### Transformation of *M* oryzae

Three-day old cultures of *M. oryzae* Guy11 or the Guy11ku80 deletion mutant, grown in 150 ml liquid complete media at 25°C and 100 rpm, were used for generation of protoplasts. The mycelia were filtered and digested with Glucanex as described above [67]. Protoplasts purification was done according to the protocol for *B. cinerea*. After washing with TMS buffer, protoplasts were suspended in TMSC buffer and adjusted to 1.5x10^8^ protoplasts per ml. For transformation, 120 µl aliquots of a protoplast suspension were mixed with the RT DNA and pre-incubated RNPs (Cas9-SV40^x4^) dissolved in 60 µl Tris-CaCl2 buffer. Then 180 µl 60% PEG 3350 were added, and the protoplast suspension was poured into CM agar containing 1.2 M sucrose for osmotic stabilization. After 24 h an upper layer containing 500 mg l^-1^ hygromycin (Hyg) or 30 mg l^-1^ Fen was poured over the agar containing the protoplasts. After 7-10 days, mutants were transferred to selection plates for further selection. RT (containing *gpd3* promotor, *hph* and *tubB* terminator) with 50 bp of homology flanks was amplified using primers MH-Alb F&R for targeting *MoAlb1*, and MH-Pit F&R for targeting *MoPIT*. For sgRNA synthesis, primers sgRNA_Alb1 and sgRNA_Pit were used. Transformants were verified using primers SeqPit F/R, SeqAlb F/R and MoPit FL F/R.

### Generation and *in vivo* selection of *sdhB* codon 272 edited strains

To be used as mixed RT for randomized editing, twenty 500 bp *sdhB* fragments differing in codon 272 (listed in Supplementary Table 4) were synthesized by Twist Bioscience (San Francisco, U.S.A.) and pool-amplified with primers TL148_ SDHB_RT_F/ TL149_ SDHB_RT_R. Illumina deep sequencing was performed to verify equal representation of each fragment (±7.5%). For PCR-based identification of edited transformants, silent mutations were introduced into the 500 bp fragments which converted an XhoI to an XbaI site (codons 278/279), and allowed differentiation between WT and edited sequences (Fig. 7a). To isolate the DNA of *sdhB* codon 272-edited transformants for sequencing, sporulation was induced on the primary transformation plates. For this, three days after transformation, the SH+Fen agar containing embedded transformants was overlaid with 0.1 volumes of 5x concentrated ME medium. After another 5-7 days, transformant conidia were harvested from densely sporulating plates. To improve the recovery of transformants, the agar discs were inverted, placed onto fresh ME (1 mg l^-1^ Fen) agar plates, and incubated again for 5-7 days until sporulation. Conidia harvested from one transformation were combined and used for DNA isolation. For sequence analysis of bulked transformants selected for resistance to SDHI, 4x10^5^ conidia of Fen^R^ transformants were inoculated in standard Petri dishes with 18 ml YBA medium (1% yeast extract, 20 g l^-1^ sodium acetate[68]) containing boscalid (0.25 mg l^-1^; BASF, Ludwigshafen, Germany), fluopyram (0.3 mg l^-1^; Bayer Crop Science, Monheim, Germany), or pydiflumetofen (0.015 mg l^-1^; Syngenta Crop Protection, Stein, Switzerland) in concentrations inhibitory for *B. cinerea* WT strain B05.10. After 72 h incubation at 20°C, conidia and germlings were harvested and used for DNA isolation [66] and sequencing (see below).

To isolate *sdhB* edited strains with defined codon 272 replacements, individual Fen^R^ transformants were purified by several transfers on ME+Fen (1 mg l^-1^), YBA+Bos (1 mg l^-1^), or YBA+Flu (2.5 mg l^-1^) agar media. Total DNA of these isolates was amplified using primers TL151_SDHB_OS_F/ TL152_SDHB_OS_R, and the 741 bp products digested with either XbaI or XhoI to test whether they were edited or WT. Edited isolates were sequenced using primer TL148_ SDHB_RT_F or TL149_ SDHB_RT_R.

### Sequencing

For deep sequencing of edited transformants, bulked *B. cinerea* DNA was first amplified in 20 µl total volume, with 2 µl DNA, 10 pM of primers sdhb_F1/ sdhb_R1, 1x MyTaq^TM^ buffer, and 1 Unit MyTaq^TM^ (Bioline; Meridian Bioscience Inc., London, UK) by incubation for 2 min at 96°C, followed by 20 cycles of 15sec 96°C, 30sec 60°C, 90sec 70°C. Nested PCR was performed in 20 µl total volume, using 2 µl of the first round PCR, under the same conditions as above, but with 15 cycles only. PCR products were purified with AmpureXP beads (Thermo Fisher Scientific, Bremen, Germany). About 100 ng of each purified PCR product was used to construct Illumina libraries using the Ovation Rapid DR Multiplex System 1-96 (NuGen Technologies, San Carlos, CA, USA). Illumina libraries were pooled and size selected by preparative gel electrophoresis. Sequencing (3 million reads per sample) was performed by LGC Genomics (Berlin, Germany) on an Illumina NextSeq 550 instrument with v2 chemistry in 2x150 bp read mode. Libraries were demultiplexed using Illumina’s bcl2fastq 2.17.1.14 software. Sequencing adapter sequences were removed from the 3’ end of reads with cutadapt (https://cutadapt.readthedocs.io/en/stable/) discarding reads shorter than 20 bp. All read pairs were filtered for valid primer combinations and reverse-complemented so that R1 corresponds to the forward primer and R2 to the reverse primer. Actual primer sequences were removed for downstream processing. Reads were quality-filtered by LGC proprietary software, removing all reads with an average Phred score below 30, and all reads containing more than 1 undetermined base (N). Subsequently, all read pairs were overlap-combined using BBMerge 34.48 from the BBMap package (https://jgi.doe.gov/data-and-tools/bbtools/bb-tools-user-guide/bbmerge-guide/). Mutated positions were identified by a custom shell script, filtering for sequences containing the motifs immediately before and after thse mutated triplet (TTTGTACAGATGT and ACTATTCTCAACTG, respectively). The sequence content between these motifs were extracted and counts for the detected sequences summarized for each sequencing library.

### Fungicide susceptibility test

Isolates with defined edits in codon 272 were tested for radial growth on YSS agar with 50 mM each of either glucose, malate, acetate or succinate [59], and for their sensitivities to SDHIs. Susceptibility to Bos (BASF), Flu (Bayer Crop Science), and Pyd (Syngenta) was assessed in the WT and in edited strains on the basis of inhibition of germination. Assays with a range of fungicide concentrations (0, 0.001, 0.003, 0.01, 0.03, 0.1, 0.3, 1, 3, 10 mg l^-1^) were carried out at 20°C. After incubation for 30 h in Greiner Bio-one polystyrene microtiter plates, the fraction of conidia containing germ tubes with lengths exceeding half of the conidial diameters was determined for each strain/fungicide pair, and an EC_50_ value (effective fungicide concentration required to inhibit germination by 50%) was calculated with the Graphpad Prism 5.01 software, using a normalized response with variable slope fitted to log fungicide concentrations.

### Microscopy

Confocal images were acquired using either a Leica SP5 (DM6000 CS), TCS acousto-optical beam splitter confocal laser scanning microscope, equipped with a Leica HCX PL APO CS 63 × 1.20. water-immersion objective or a Zeiss LSM880, AxioObserver SP7 confocal laser-scanning microscope, equipped with a Zeiss C-Apochromat 40x/1.2 W AutoCorr M27 water-immersion objective. Fluorescence signals of GFP (Leica: excitation/emission 488 nm/500-550 nm, Zeiss: excitation/emission 488 nm/500-571 nm), were processed using Leica software LAS AF 3.1, Zeiss software ZEN 2.3 or Fiji software.

### Protein analysis

For in-gel detection of superoxide dismutase activity, *B. cinerea* conidia were germinated in ME medium overnight, washed with extraction buffer (100 mM potassium phosphate buffer (pH 7.8) 0.1 mM (EDTA) 1 % (w/v) polyvinyl-pyrrolidone (PVP) 0.5% (v/v) Triton X 100) the mycelium ground with mortar and pestle in liquid nitrogen. Fifteen µg of cleared extract was separated in an polyacrylamide gel and stained for SOD activity as described [69]. For detection of Cas9 Sod1-GFP fusion proteins, *B. cinerea* protein extracts prepared as described above were separated in an SDS polyacrylamide gel and subjected to an immunoblot on nitrocellulose, using monoclonal antibodies against Cas9 (Clontech, Palo Alto, CA, USA) or GFP (Sigma), followed by chemiluminescent detection.

### Statistics and reproducibility

Statistical analyses were carried out with the GraphPad Prism software. The detailed analysis method is depicted in the individual figure legends. All experiments were carried out at least three times. For growth and infection assays, three technical replicates per sample were performed. Box limits of box plots represent 25th percentile and 75th percentile, horizontal line represents median. Whiskers display minimum to maximum values. Bar charts represent mean values with standard deviations.

## Acknowledgements

We are grateful to Sabine Fillinger (INRA, Paris, France) for providing telomere plasmid pFB2N, and to Pinkuan Zhu (Shanghai Normal University, China) for help with pTEL constructions. We thank Patrick Pattar for excellent technical support, and Nora Fischbach for help with characterization of edited strains. Andrew Foster is kindly acknowledged for providing Guy11ku80. This work was supported by BioComp initiative of Rhineland-Palatinate. Alex Wegner was supported by a PhD grant of RWTH Aachen University.

## Supplementary Figures and Tables

**S1 Fig.**
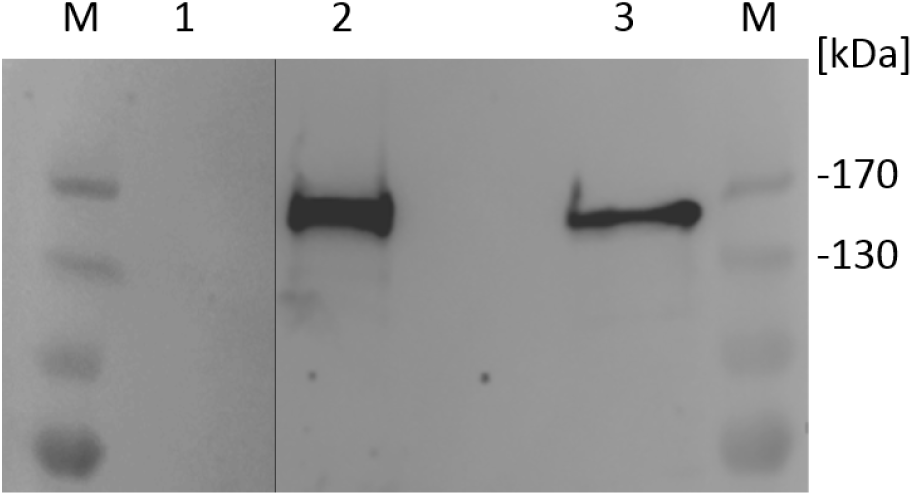
Detection of Cas9 expression in *B. cinerea*. Total *B. cinereal* protein extracts (15 µg per lane) were loaded, separated by polyacrylamide gel electrophoresis, and Cas9 detected with a monoclonal Cas9 antibody. M: Marker; 1: B05.10 (WT); 2: B05.10-Cas9-SV40^x4^; 3 B05.10 (pTEL-Cas9-Stu^x2^).

**S2 Fig.**
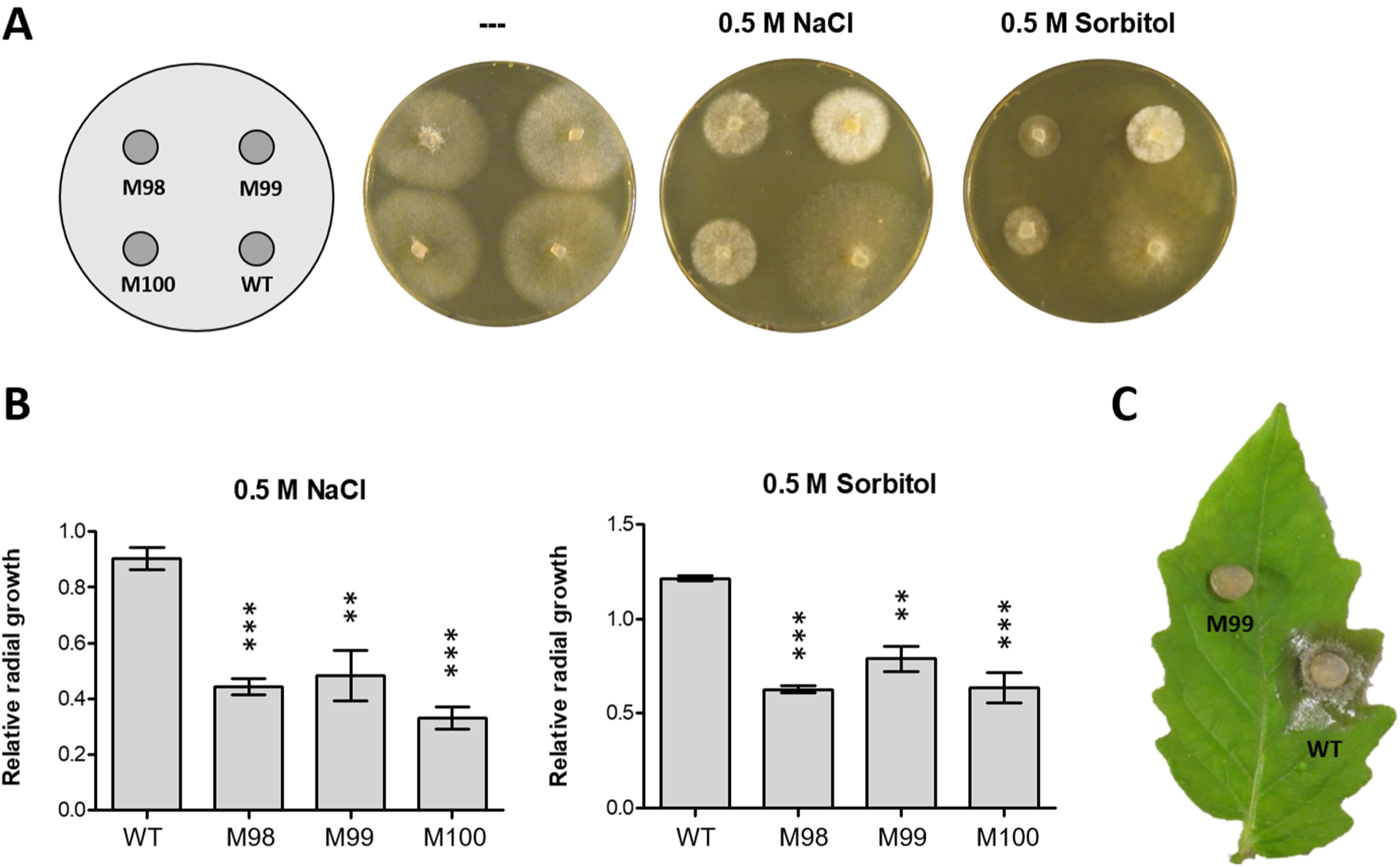
Sensitivity to osmotic and salt stress and virulence of *B. cinerea* WT and CRISPR/Cas-induced *Bos1* mutants. (A) Pictures of three Ipr^R^ *Bos1* mutants (M98, M99, M100, all having the same ‘+T’ insertion) and WT growth for 48 h on ME medium without (---) and with 0.5 M NaCl or sorbitol. (B) Effects of salt and osmotic stress treatments on radial growth, compared to growth on pure ME medium (n=3). The *p* values by one-way ANOVA followed by Dunnett’s multiple comparisons (control: WT) post hoc test are indicated. \*\**p* ≤ 0.01; \*\*\**p* ≤ 0.001. (C) Infection on tomato leaf by WT and mutant M99 (72 h).

**S3 Fig.**
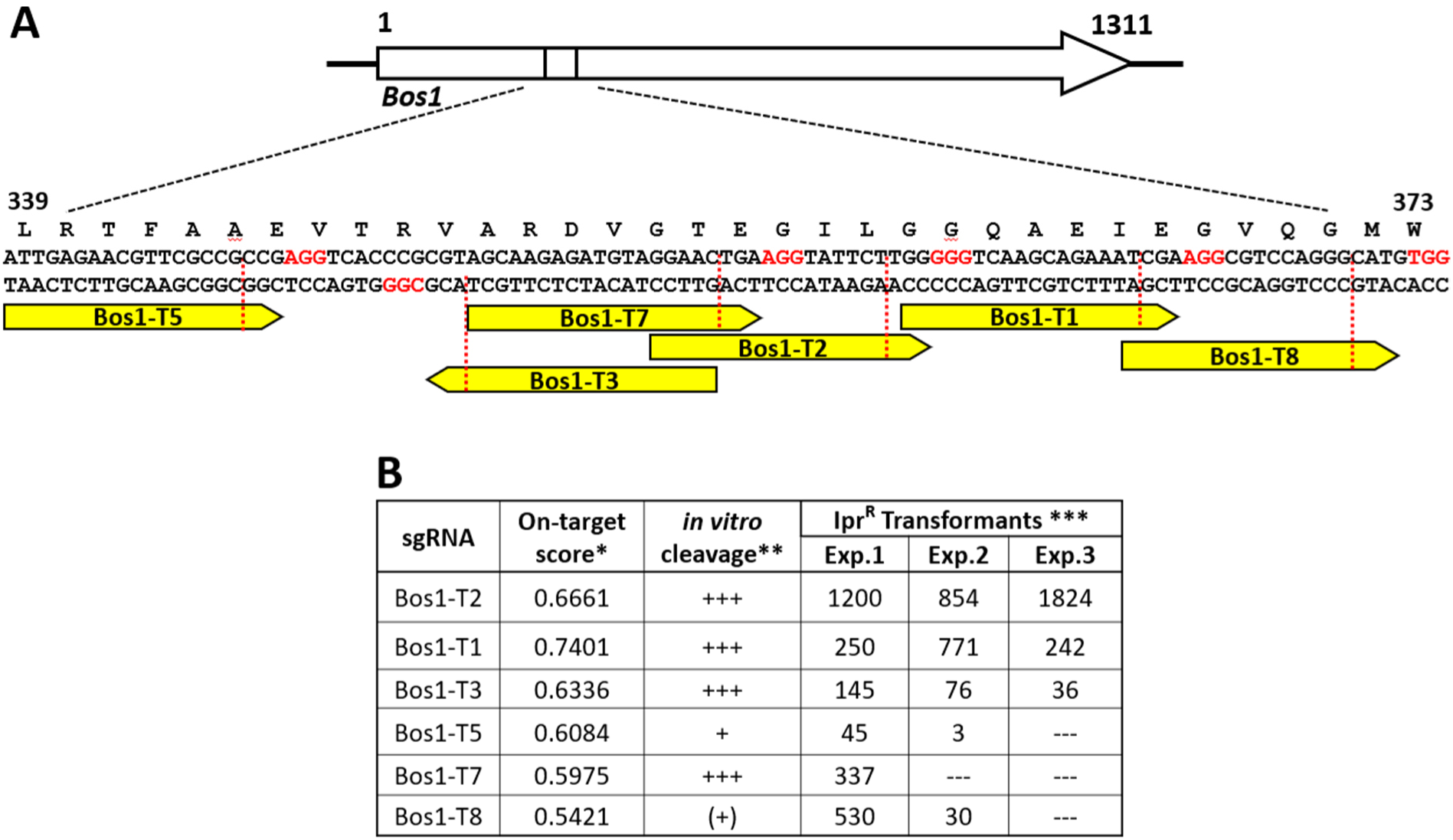
*In vitro* and *in vivo* CRISPR/Cas performance of different sgRNAs targeting *Bos1*. (A) Positions and expected cleavage sites (red dotted lines) of the sgRNAs (yellow) in *Bos1*. PAM sequences for each of the sgRNAs are indicated in red. (B) Summary of on-target scores, *in vitro* cleavage activities, and transformation efficiencies with different sgRNAs. *On-target efficiency calculated with the Broad Institute GPP sgRNA Designer. **Estimated from gel pictures. *** Number of Ipr^R^ *B. cinerea* mutants per assay.

**S4 Fig.**
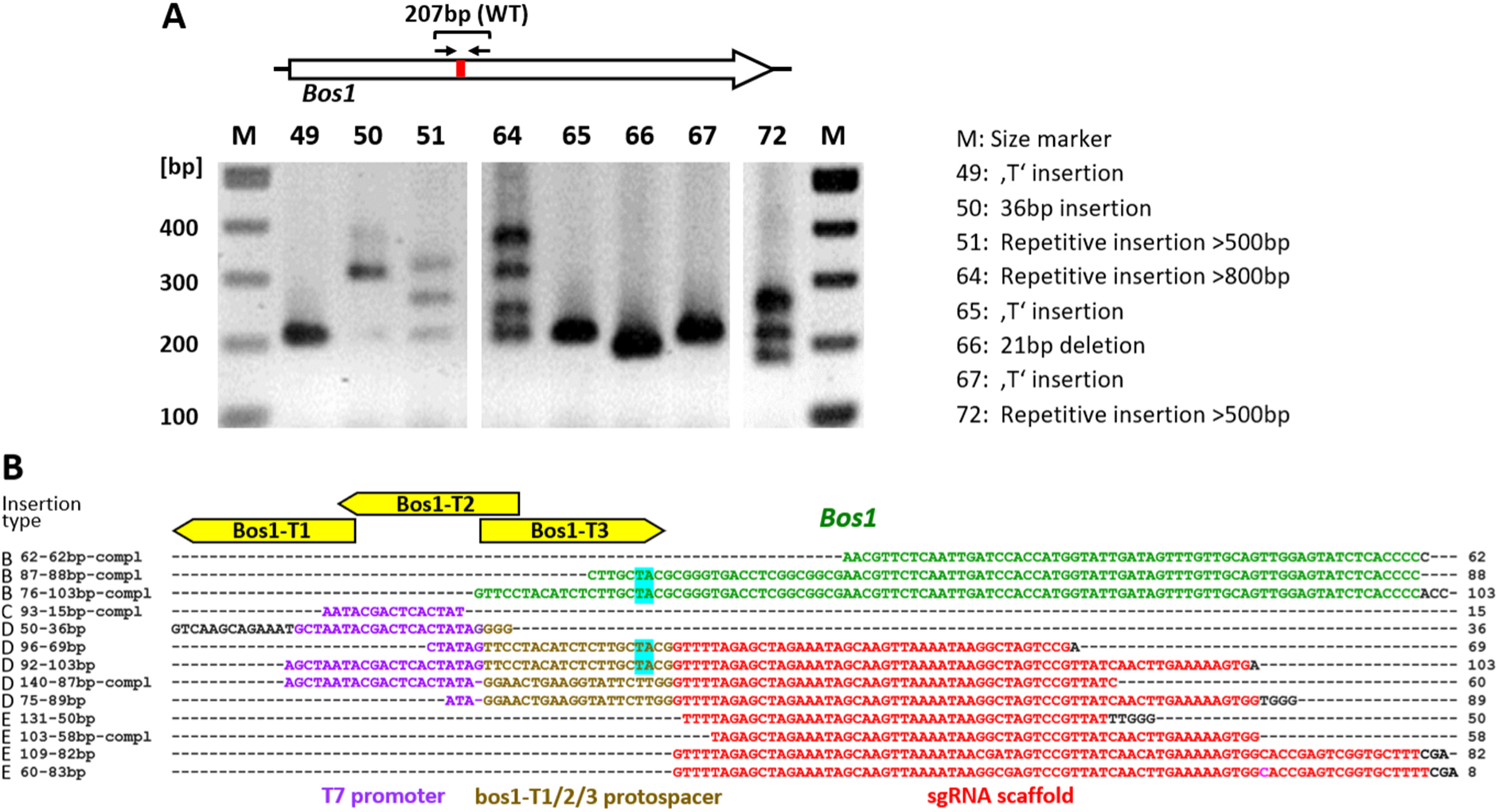
Analysis of PCR fragments covering CRISPR/Cas-induced cleavage-repair sites in *Bos1*, from Ipr^R^ transformants. (A) Stained agarose gel with PCR fragments generated with primers TL_87 Bos1_check 200 Fw/ TL_87 Bos1_check 200 Rv, showing variations of fragment sizes due to different types of NHEJ-induced mutations in transformants obtained with Cas9/bos1-T1 RNP. (B) Origins and sequences of different NHEJ insertion types obtained with different Cas9/bos1 RNPs. Type A (not shown): 164 bp *B. cinerea* mitochondrial DNA, two joined fragments of 84 and 79 bp. Type B: *B. cinerea Bos1*-DNA. Type C: 15 bp fragment of the sgRNA scaffold encoding part of the T7 RNA polymerase promoter. Type D: sgRNA scaffold DNA containing part or all of the protospacer sequences of bos1-T1/-2/-3. Type E: sgRNA scaffold DNA lacking protospacer sequences.

**S5 Fig.**
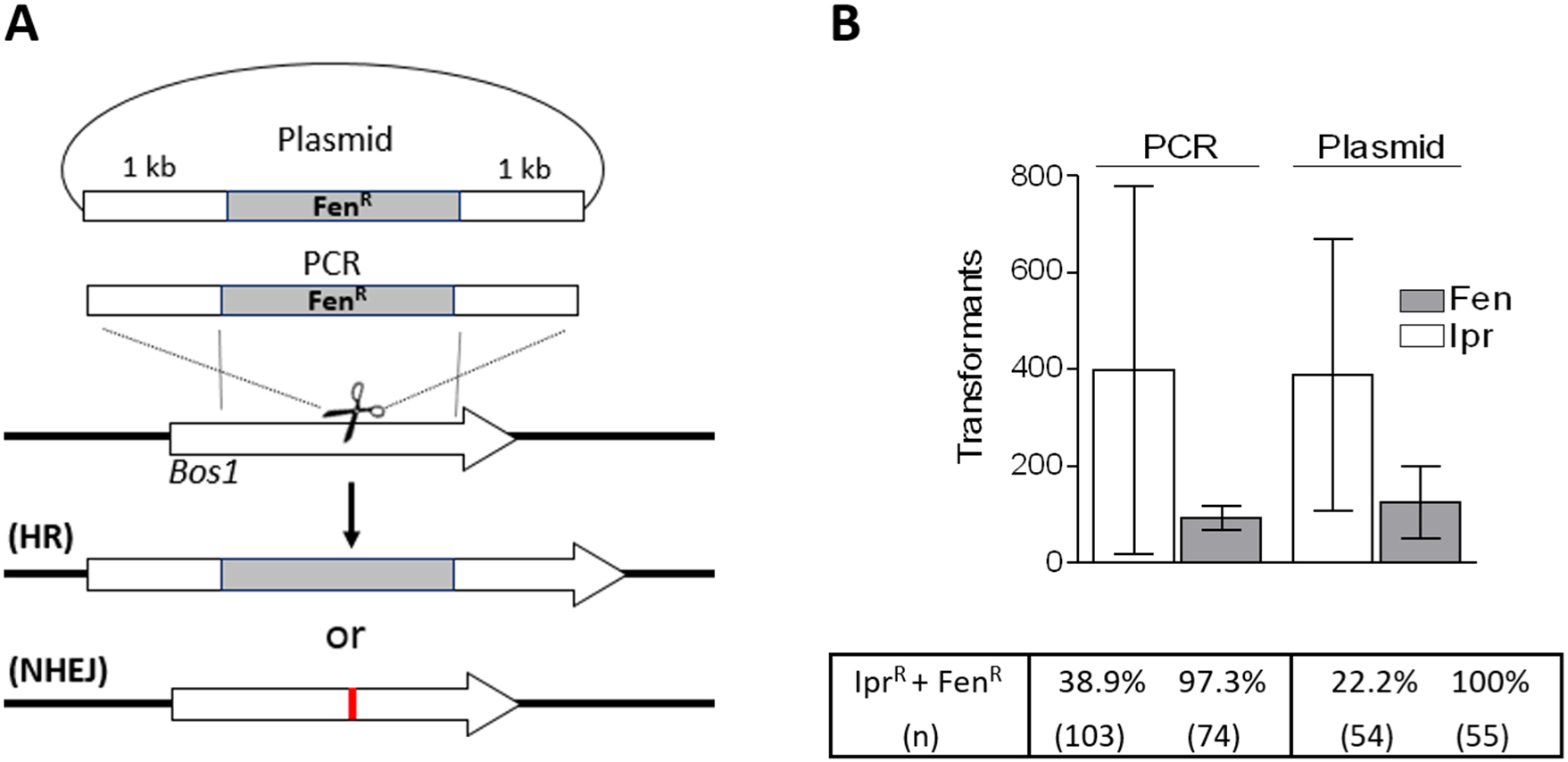
Transformation of Cas9/*Bos1*-T2B-gRNA RNP and Fen^R^ RT with 1 kb *Bos1* homology flanks. (A) Experimental scheme. *Bos1* inactivation leading to Ipr^R^ occurs either by targeted integration of the Fen^R^ RT via HR, or via NHEJ. (B) Transformation results: Primary selection was either for Ipr^R^ (white bars) or for Fen^R^ (grey bars) (n=3). Below the diagram, the fraction of transformants with resistance to both fungicides is shown. (n): Number of transformants tested. Statistical analyses were performed by analysis of variance (ANOVA, followed by Dunnett’s multiple comparisons. No significant differences between transformation results with PCR fragments and circular plasmids, or between Ipr^R^ and Fen^R^ colony numbers were observed.

**S6 Fig.**
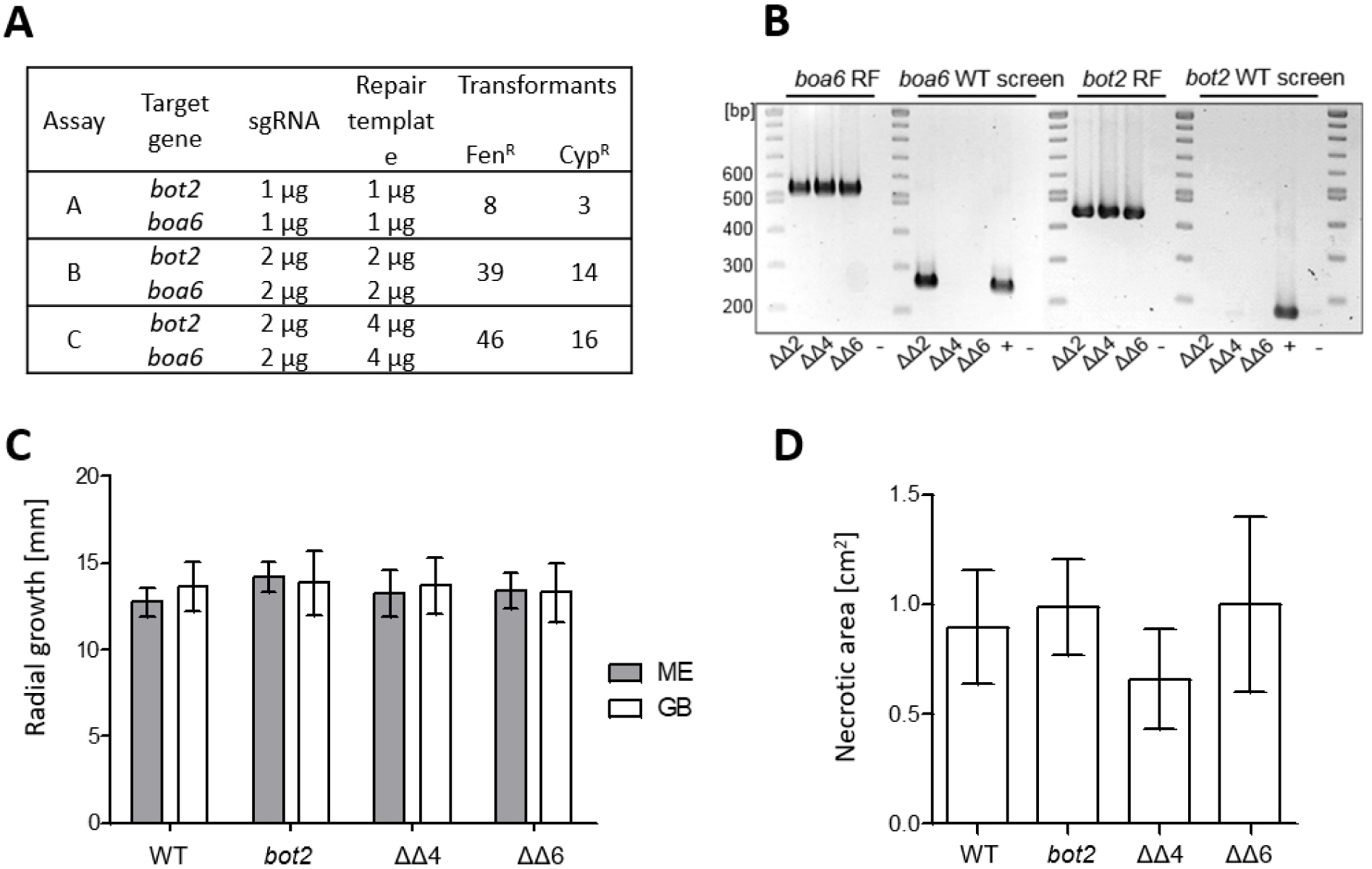
CRISPR/Cas-HR-mediated single and double k.o. mutagenesis of *bot2* and *boa6*. (A) Transformation results. (B) PCR-based verification of *bot2 boa6* double (ΔΔ) k.o. mutants. *boa6* RT right flank (RF) integration screen using primers TniaD_ol_Cyp_Fw/TL129 (537 bp); *boa6* WT screen using primers TL157/TL158 (263 bp), *bot2* RT RF integration screen using primers TL130/TL132 (444 bp), *bot2* WT screen using primers TL133/TL159 (180 bp). (C) Growth of WT and mutants after 72 h on agar plates with rich (ME) and minimal (GB: Gamborg GB5 with 25 mM glucose) medium (one way ANOVA; n=3). (D) Lesion formation after 72 h on tomato leaves (one way ANOVA; n=3). In (C) and (D), no significant differences in radial growth and infection between WT and mutants were observed.

**S7 Fig.**
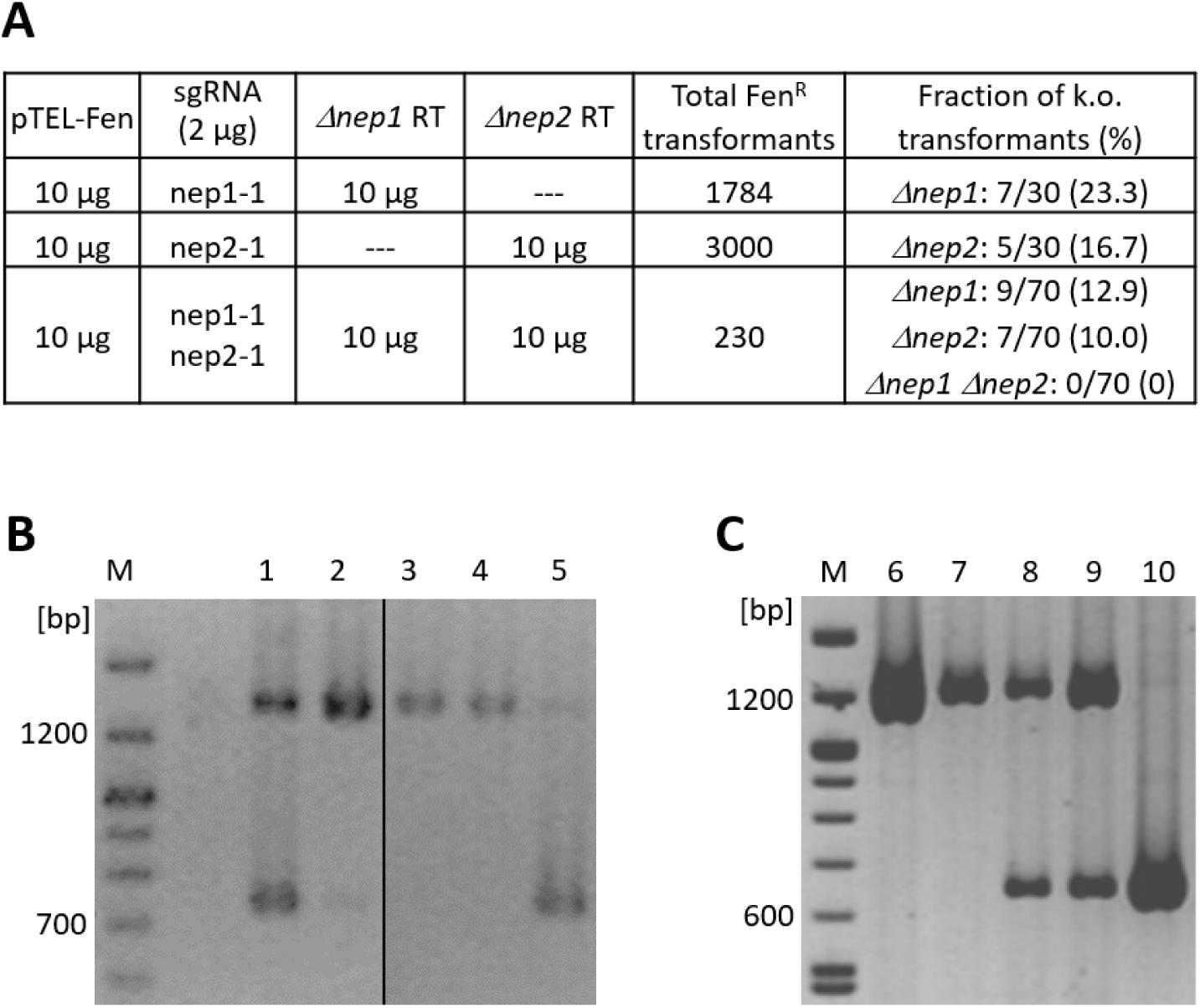
Use of pTEL-Fen for marker-free k.o. mutagenesis of *nep1* and *nep2*. (A) Transformation result. (B) PCR-based verification of *nep1* deletion mutants, using primers TL143/TL144; size of WT fragment 1,353 bp, size of *nep1* k.o. fragment 733 bp. Lanes 1-5: Fen^R^ transformants. Transformant #5 represents a nearly pure *nep1* mutant. (C) PCR-based verification of *nep2* deletion mutants, using primers TL145/TL146; size of WT fragment 1,220 bp, size of *nep2* k.o. fragment 641 bp. Lane 6: *B. cinerea* WT; lanes 7-10: Fen^R^ transformants. Transformant #10 represents a purified *nep2* mutant. M: DNA marker.

**S8 Fig.**
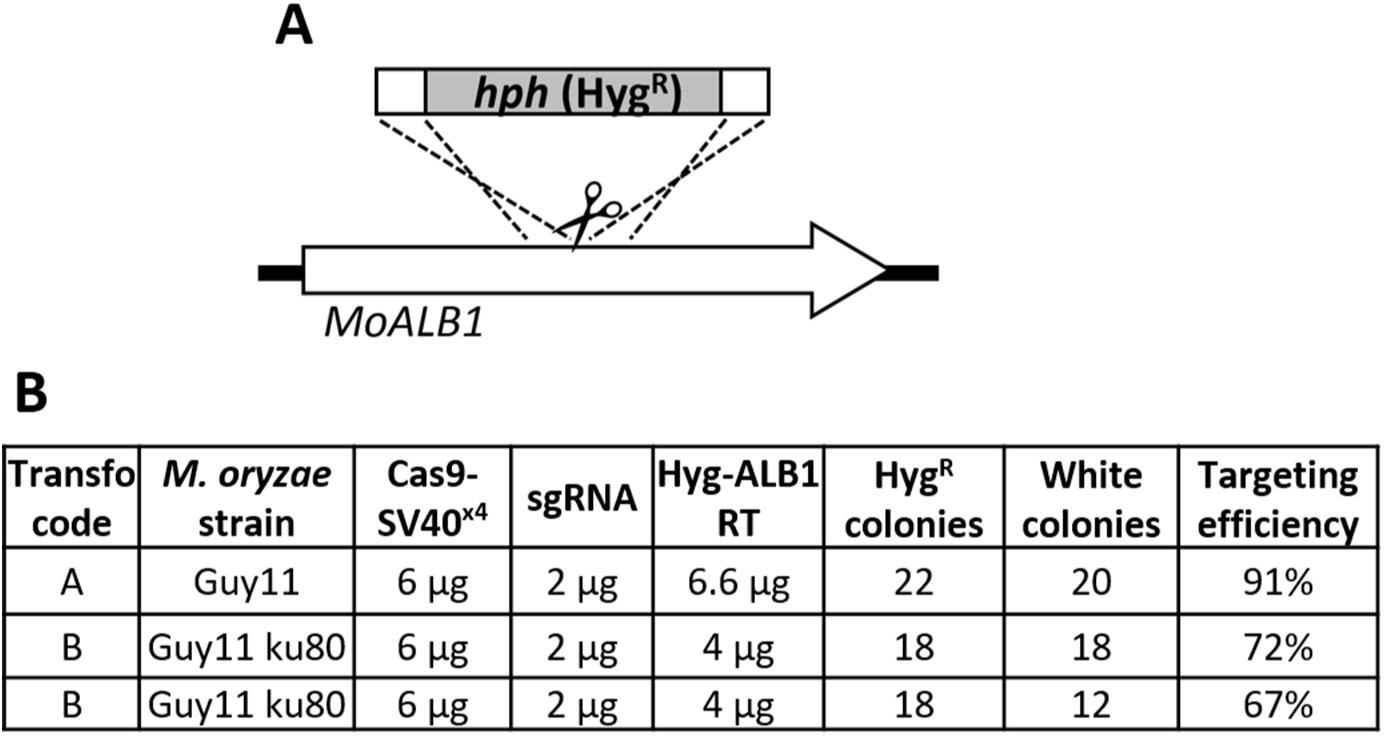
CRISPR/Cas efficiency with RNP in *M. oryzae*. (A) Scheme of CRISPR/Cas targeting of *MoALB1*, using a repair template with a hygromycin resistance cassette and 50 bp homology flanks. (B) CRISPR/Cas components used and results of transformations with *M. oryzae* strain Guy11 and Guy11 ku80.

**S9 Fig.**
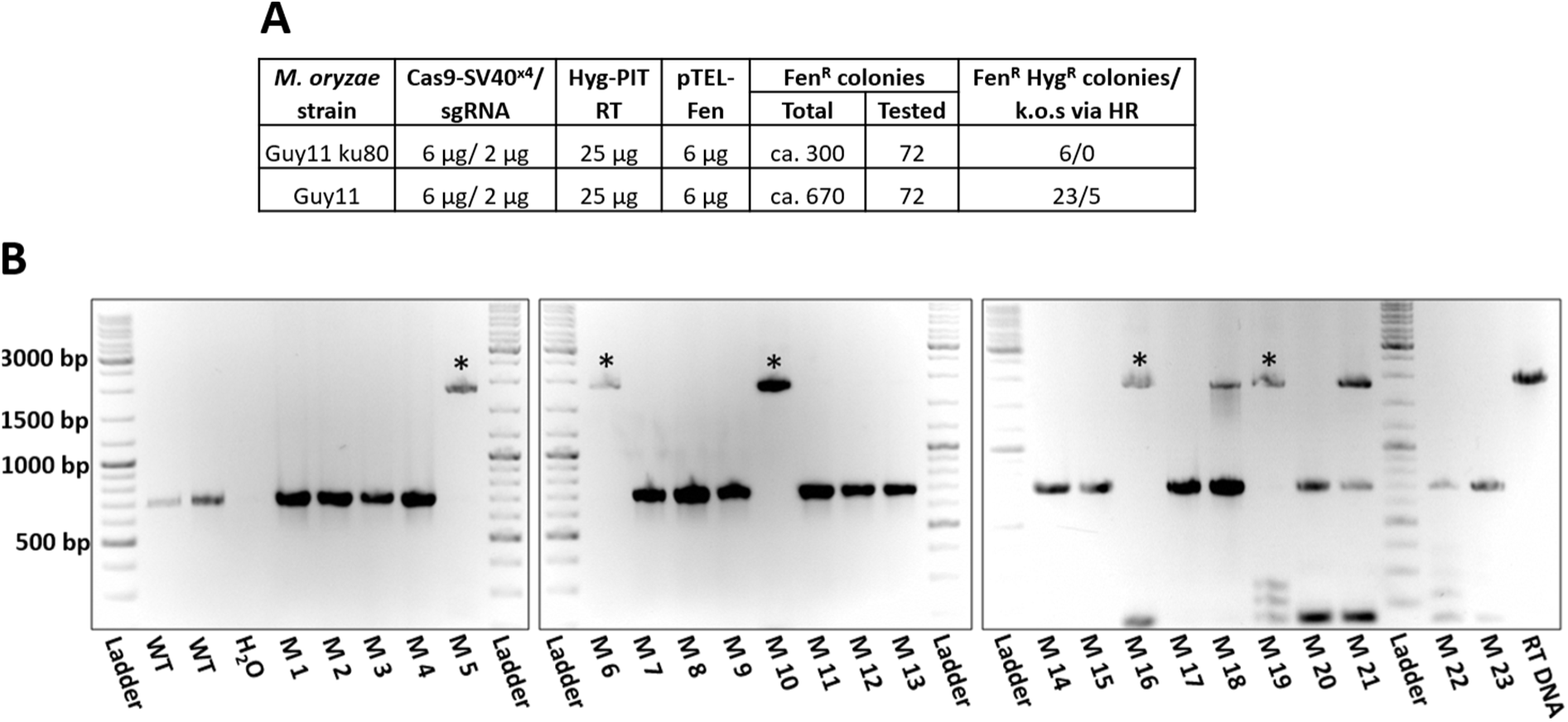
pTEL-mediated k.o. of *MoPIT* via HR in *M. oryzae*. (A) Results of transformations with strains Guy11 and Guy11ku80. (B) PCR-based identification of strain Guy11 *MoPIT* k.o. mutants generated by coediting, using primers MoPit FL F and MoPit FL R. HR was confirmed by sequencing PCR product amplified with primers SeqPit_F/ SeqPit_R.

**S10 Fig.**
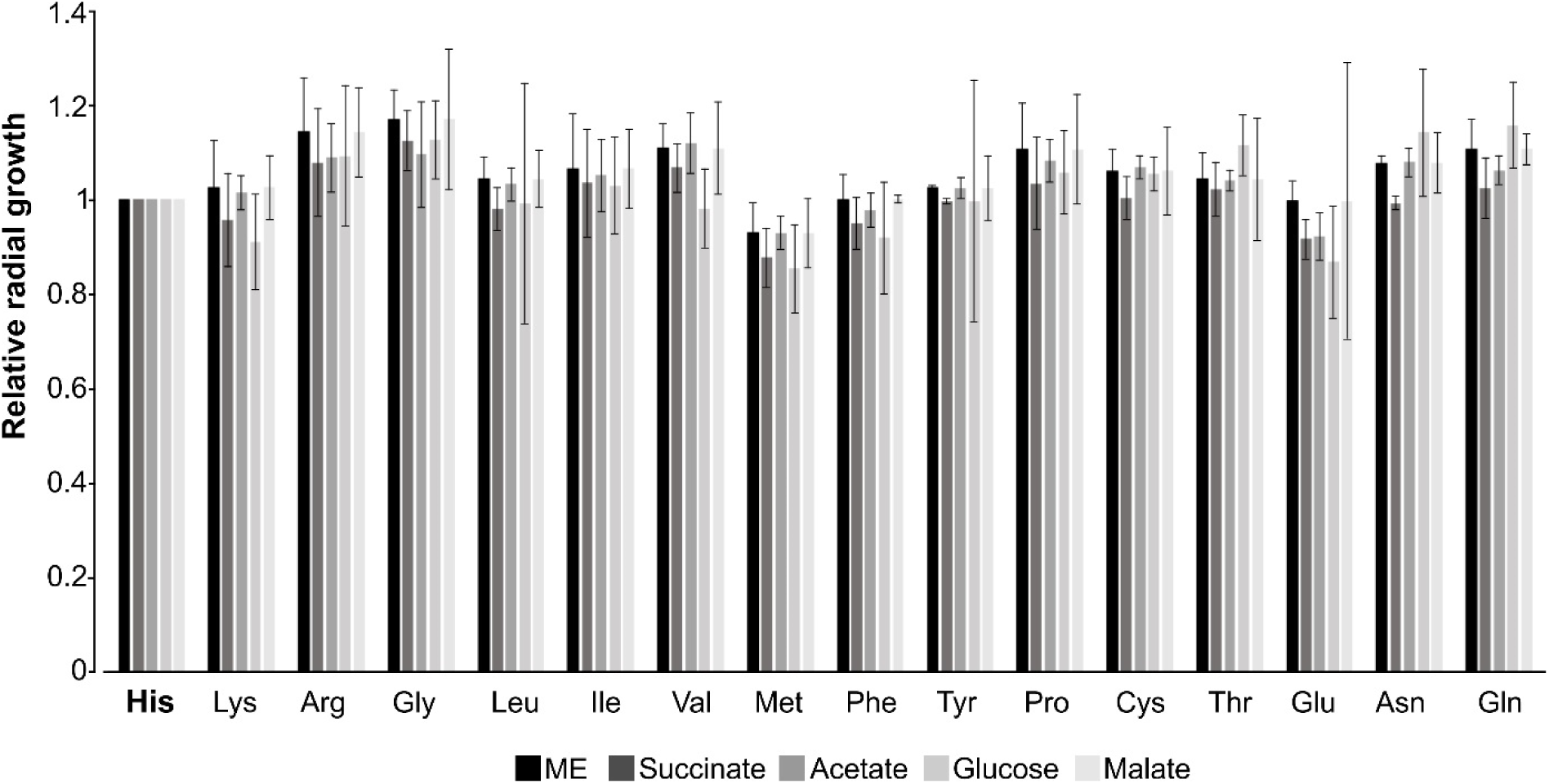
Mycelium growth after 72 h of *B. cinerea sdhB* codon 272 exchange mutants on agar media containing ME or YSS with different carbon sources (50 mM each), relative to the WT strain (n=3). Statistical analyses were performed by analysis of variance (ANOVA) followed by Dunnett’s multiple comparisons (control: His). No significant differences between the growth rates of the WT strain (His) and any of the mutants were observed.

**S1 Table.** pTEL-mediated coediting via NHEJ in *M. oryzae*. CRISPR/Cas components used and results of (co-) transformations with strain Guy11 and Guy11 ku80. Transformations with the same letter were done with the same batch of protoplasts.

| Experiment | *M. oryzae* strain | Cas9-SV40x4/<br>sgRNA amounts | pTEL-Fen |  |  | White Hyg<sup>R</sup> colonies<br>(coediting rate) |
| --- | --- | --- | --- | --- | --- | --- |
|  |  |  |  | Total | Tested |  |
| B | Guy11 | --- | 1 µg | 18 | --- | --- |
| B | Guy11 | --- | 2 µg | 200 | --- | --- |
| C | Guy11 | --- | 2 µg | 555 | --- | --- |
| C | Guy11 | --- | 4 µg | 3,800 | --- | --- |
| D | Guy11 | --- | 4 µg | 4,000 | --- | --- |
| D | Guy11 ku80 | --- | 4 µg | 3,000 | --- | --- |
| E | Guy11 ku80 | --- | 12 µg | 1,500 | --- | --- |
| E | Guy11 ku80 | 6 µg/ 2 µg | 12 µg | 300 | 72 | 5 (7%) |
| B | Guy11 | 6 µg/ 2 µg | 6 µg | 96 | 36 | 17 (47%) |
| C | Guy11 | 6 µg/ 2 µg | 6 µg | 280 | 35 | 17 (49%) |
| C | Guy11 | 6 µg/ 2 µg | 3 µg | 120 | 36 | 13 (36%) |

**S2 Table.** Results of cotransformation of *B. cinere*a with pTEL-Fen, sdhB272-sgRNA-RNP and 500 bp sdhB-272 (x20) repair template.

| Transformation<sup>1</sup> | sgRNA | Fen<sup>R</sup> colonies/<br>transformation | Sequenced<br>transformants | Editing frequency |  |
| --- | --- | --- | --- | --- | --- |
|  |  |  |  | Counted<sup>2</sup> | Illumina seq. |
| A | sdhB272-1 | 820 | --- | 9/72 (12.5%) | n.a. |
| A | sdhB272-2 | 1,100 | --- | 12/96 (12.5%) | n.a. |
| B | sdhB272-1 | 2,680 | --- | 10/41 (24%) | n.a. |
| C | sdhB272-2 | 2,200 | --- | 11/26 (41%) | n.a. |
| D | sdhB272-2 | 4,820 | 6,100 | n.a. | 31.6 |
| E | sdhB272-2 | 6,680 | 6,200 | n.a. | 21.4 |
| F | sdhB272-1 | 7,080 | 10,400 | n.a. | 11.4 |

<sup>1</sup> 2x10<sup>7</sup> protoplasts were transformed with 10µg pTEL-Fen, 6µg Cas9-2µg sdhB272-RNP, and 10µg sdhB-272(x20) repair template. <sup>2</sup>To determine the fraction of edited transformants, DNA of individual transformants was prepared, amplified with primers TL151\_SdhB\_OS\_F/ TL152\_SdhB\_OS\_R covering the edited region, and digested with either XhoI (cut in WT *sdhB* DNA) or XbaI (cut in edited *sdhB* DNA). n.a.: Not analysed.

**S3 Table.**
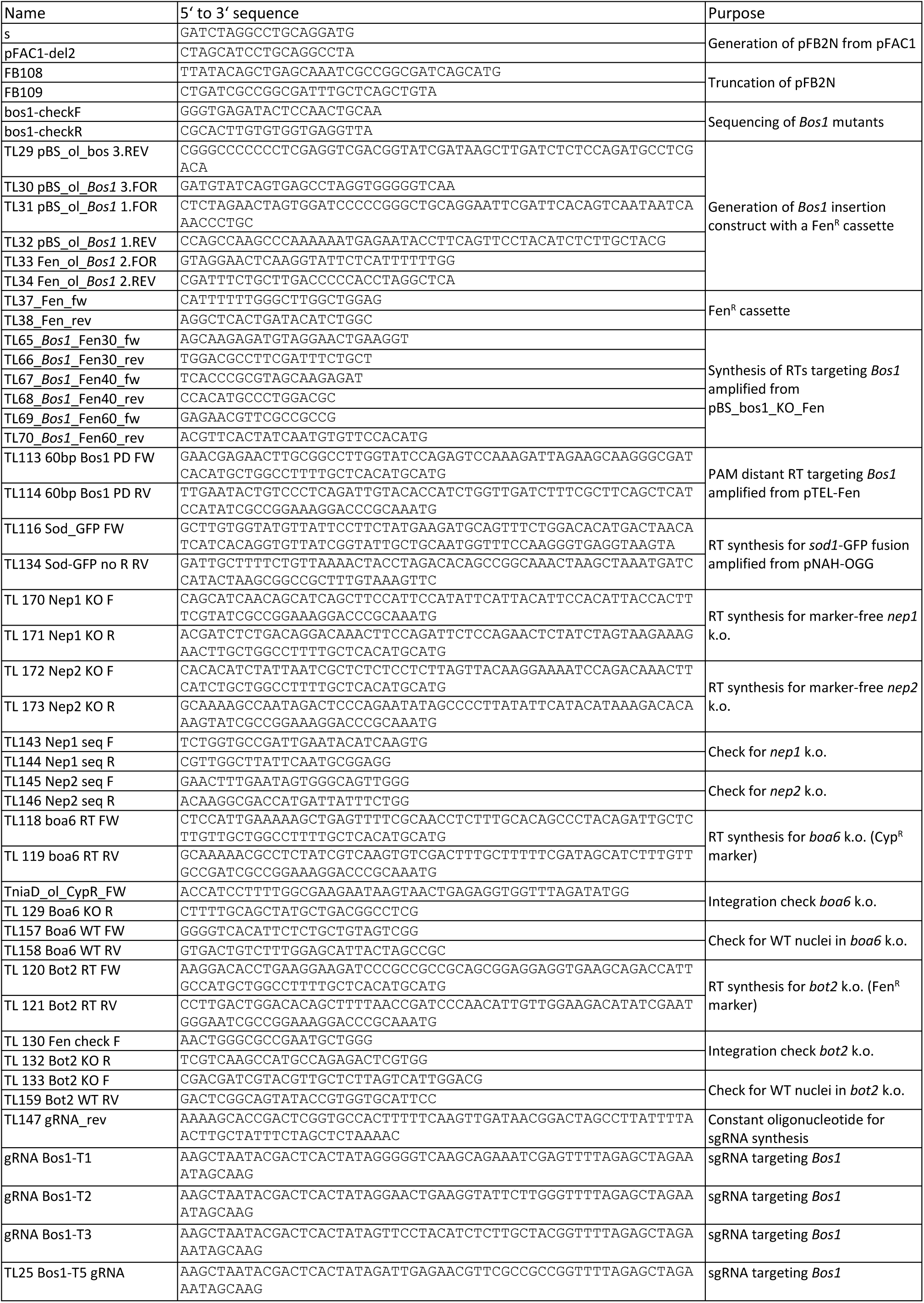
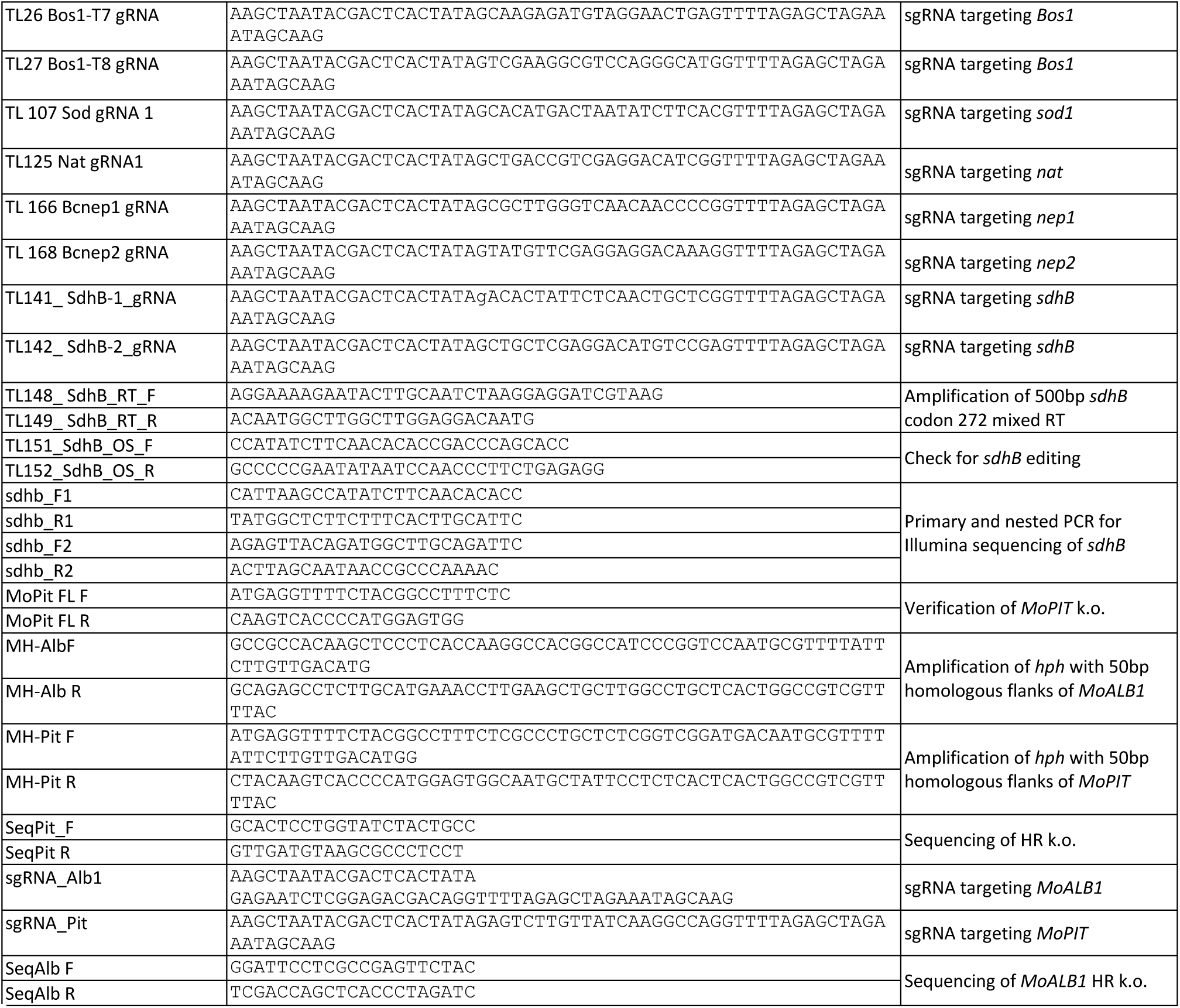
Oligonucleotides used.

